# Mesoscale proximity labeling to study macro changes to chromatin occupancy

**DOI:** 10.1101/2025.03.13.643041

**Authors:** Cameron J. Douglas, Preston Samowitz, Feifei Tong, Alice Long, Caitlin M. Bradley, Laszlo Radnai, David W. C. MacMillan, Courtney A. Miller, Gavin Rumbaugh, Ciaran P. Seath

## Abstract

Proximity labeling traditionally identifies interactomes of a single protein or RNA, though this approach limits mechanistic understanding of biomolecules functioning within complex systems. Here, we demonstrate a strategy for deciphering ligand-induced changes to global biomolecular interactions by enabling proximity labelling at the mesoscale, across an entire cellular system. By inserting nanoscale proximity labelling catalysts throughout chromatin, this system, MesoMap, provided new insights into how HDAC inhibitors regulate gene expression. Furthermore, it revealed that the orphaned drug candidate, SR-1815, regulates disease-linked *Syngap1* gene expression through direct inhibition of kinases implicated in both neurological disorders and cancer. Through precise mapping of global chromatin mobility, MesoMap promotes insights into how drug-like chemical probes induce transcriptional dynamics within healthy and disease-associated cellular states.

## Introduction

Changes in chromatin architecture promote gene expression dynamics arising from intrinsic cellular regulation and adaptive responses initiated by external stimuli. Many methods that interrogate changes in chromatin structure have emerged and they have become mainstay techniques in molecular and cellular biology for studying genomic accessibility (ATAC-seq), or genome wide looping interactions between sites in the genome (Hi-C) *(1, 2)*. However, these methods are based upon nucleic acid sequencing and can only infer information about protein localization. Chromatin immunoprecipitation methods enable visualization of genomic localization changes for a single target protein, but is limited by antibody availability, and can be cost prohibitive to study more than a few proteins at a time.

Proximity labeling (PL) is a powerful method for identifying protein-protein interactions in a cellular context, particularly for the study of insoluble proteins and those scaffolded by lipids and nucleic acids. Generally, PL approaches are utilized to study the interactome of a single protein, and how said protein responds to external stimuli, changes in protein structure, or localization *(3, 4)*. Despite significant advances, interactomics methods are of limited use when assessing a stimulus with an unknown mechanism, such as bioactive ligands derived from a phenotypic screen.

A method that captures protein occupancy dynamics across the entire face of chromatin would provide mechanistic insights into how bioactive ligands alter transcriptional states. To develop such a technique, two specific challenges must be addressed. First, the commonly used PL methods (APEX2, BioID) are unable to differentiate between the chromatin environment and the bulk nuclear proteome due to long labeling radii, although they have been used effectively to study translocation of proteins and RNA from one organelle to another *(5, 6)*. This impedes distinguishing chromatin bound proteins from nuclear localized proteins, as both would be labeled by these existing methods. Second, no single protein PL target can encompass both the entirety of the chromatin microenvironment and the heterogeneity of chromatin structure.

We recently reported a nanoscale PL method to study histone interactomes that utilizes the µMap photo-proximity labeling platform, which only labels proteins directly bound to chromatin, rather than the entire nucleus *(7)*. Based on that precedent, we questioned whether µMap could be adapted to study protein occupancy across the entirety of chromatin, an approach we term mesoscale proximity labeling, or MesoMap. In principle, this could be accomplished by incorporating µMap photo-proximity labeling catalysts throughout chromatin using multiple baits, which would enable measurements of changes across the entirety of chromatin, rather than at a specific nuclear protein **(Fig. 1A)**. If successful, this strategy would provide an unbiased chromatin-wide readout of how a cellular stimulus changes molecular interactions that lead to transcriptional phenotypes.

**Fig. 1.**
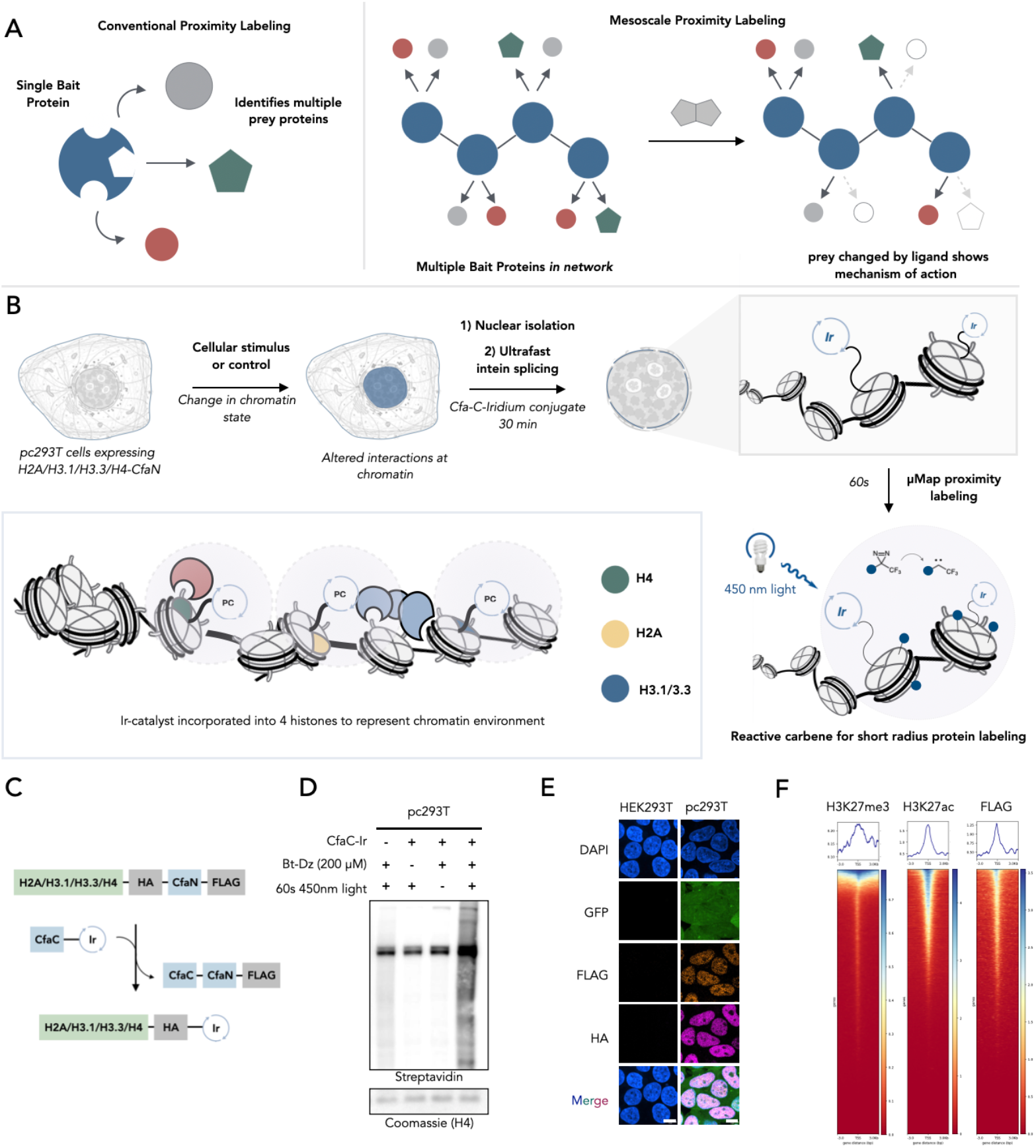
Development of the mesoscale proximity labeling platform MesoMap. (**A**) Traditional proximity labeling is used to map the interactome of a single bait protein. Mesoscale proximity labeling implements multiple bait proteins to capture system wide changes to protein interactions in response to small molecule stimuli (**B**) Scheme illustrating workflow of MesoMap proximity labeling in which biochemically intact nuclei are isolated, CfaC-Ir photocatalyst is spliced onto histone constructs, proximity labeling is conducted with biotin-diazirine probes and blue light irradiation, and protein interactions along the surface of chromatin are captured. (**C**) Scheme of ultrafast intein splicing to append an iridium catalyst to histone constructs. (**D**) Western blot of pc293T cells, showing labeling only when pc293T are spliced with CfaC-Ir, incubated with Bt-Diazirine, and irradiated for 1 min with 450 nm light. (**E**) Immunocytochemistry of pc293T cells stained for HA and FLAG. GFP signal is free GFP reporter of construct. DNA is stained with DAPI. Scale bar = 10 µm. (**F**) Heatmaps of CUT&Tag of pc293T cells showing H3K27me3, H3K27ac, and FLAG peaks centered on transcription start site +/-3 kb. Representative track of one of three biological replicates. Reads normalize to counts per million. Generated using deepTools.

## Results

### Methodology development

We generated a HEK293T cell line stably expressing four histones bearing CfaN fusions in addition to free GFP for selection: H2A, H3.1, H3.3, and H4. H2A and H4 provide spatial differences within a single nucleosome, while H3.1 and H3.3 are incorporated into different regions of chromatin. This strategy should provide a reasonable distribution of photocatalysts throughout the entirety of chromatin and they would be positioned to label only chromatin bound proteins through nanoscale photo-proximity labeling **(Fig. 1B)**. To incorporate the histones uniformly, we utilized the MultiMam expression cassette bearing all of the CfaN tagged histones and propagated single cell clones *(8)*. After expansion, colonies underwent flow cytometry to select a cell line with uniform, high expression of GFP (**Fig. S1A**), referred to as pc293T herein. Isolation of biochemically intact nuclei from pc293T cells enabled rapid intein splicing using CfaC-biotin (**Fig. S1B**), in addition to the iridium photocatalyst conjugated CfaC after 30 mins (**Fig. 1C and Fig. S1C**). Catalyst conjugated chromatin was shown to undergo photocatalytic proximity labeling in a light and probe dependent manner **(Fig. 1D)**. Confocal microscopy of pc293T cells showed exclusive nuclear localization of the fluorescently labeled histone constructs **(Fig. 1E)**. Distribution of the transgenes throughout chromatin was validated by CUT&TAG sequencing using anti-FLAG antibodies and showed distribution of the construct throughout heterochromatin (H3K27me3) and euchromatin (H3K27ac) (**Fig. 1F** and **Fig. S1D**).

Chemoproteomics was conducted to validate that our method provides a unique chromatin interactome rather than capturing the bulk nuclear proteome. We irradiated biochemically intact nuclei containing catalyst conjugated chromatin in the presence of 200 µM diazirine-biotin for 60 seconds. Excess diazirine-biotin was washed out prior to lysis and streptavidin enrichment. We compared the nuclear proteome with labeled proteins from our mesoscale protocol. Here, the MesoMap protocol provides a distinct interactome from the nuclear proteome (**Fig. S1E** and **Table S1**). We identified several epigenetic and chromatin remodeling complexes, in addition to proteins associated with both heterochromatin and euchromatin (**Fig. S1F**). GO analysis of the proteins enriched in our PL samples also showed enrichment of chromatin related terms such as chromatin remodeling and mitotic cell cycle (**Fig. S1G**). Conversely, GO terms associated with RNA metabolism were enriched in the bulk nuclear samples, supporting our hypothesis that our carbene based proximity labeling protocol cannot diffuse through the bulk nuclear milieu (**Fig. S1G**).

### The impact of HDAC inhibitors on chromatin occupancy of epigenetic complexes

We next sought to test our method by measuring chromatin state changes induced by treatment with the histone deacetylase inhibitor, romidepsin **(Fig. 2A-B)**. Romidepsin promotes histone acetylation by inhibiting multiple deacetylase enzymes, leading to a more open chromatin state and altered transcriptional signatures *(9)*. This mechanistic paradigm has been proposed as a treatment for many indications, including neurological diseases, several cancers, and HIV *(9*–*11)*.

**Fig. 2.**
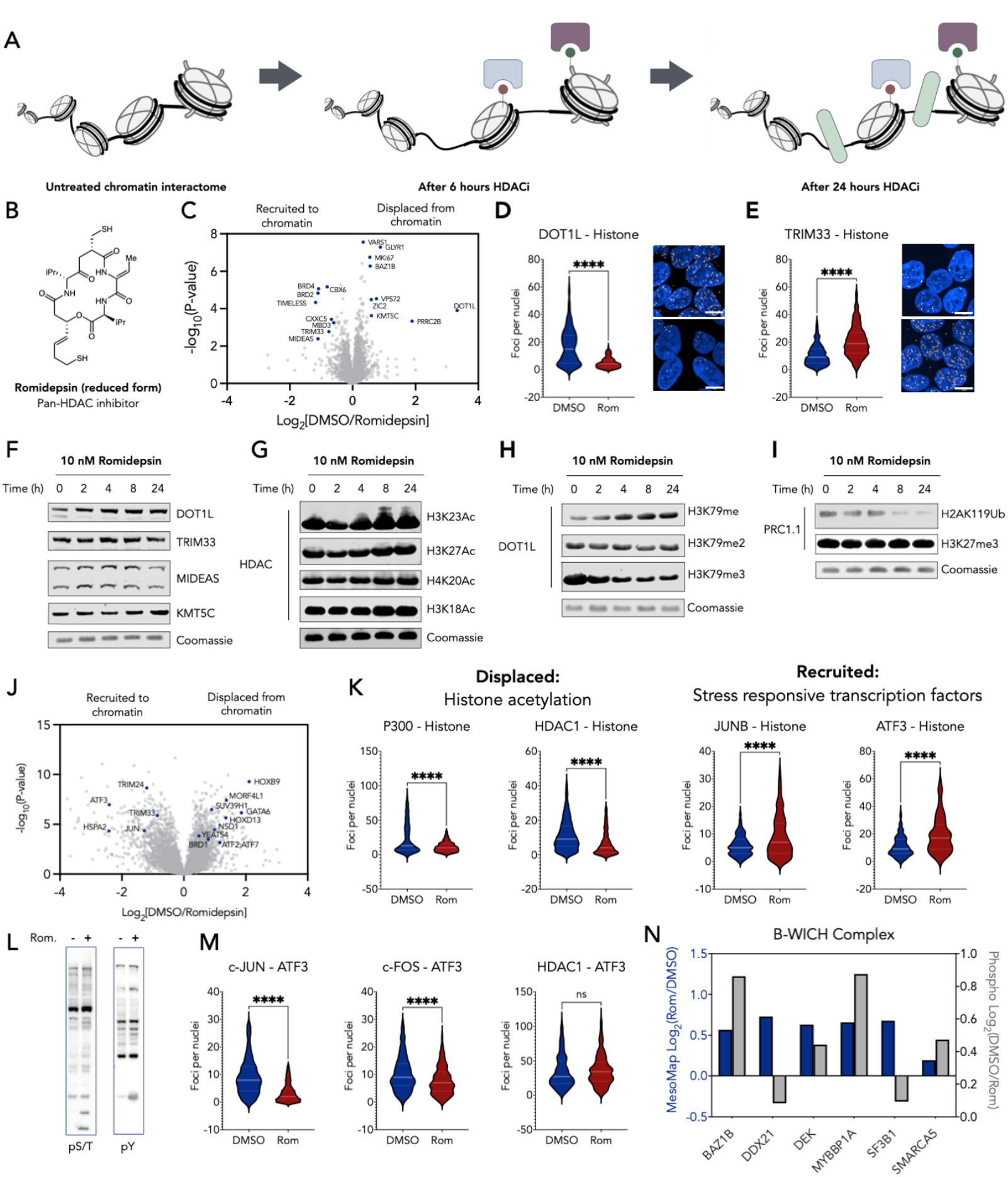
MesoMap investigation of HDAC inhibitor effect on chromatin interactome. (**A**) Graphic showing changes to chromatin over time in response to treatment with 10 nM romidepsin (**B**) HDAC inhibitor romidespin. (**C**) Volcano plot of proximity proteomics of pc293T cells treated with either 10 nM romidepsin or DMSO for 6 h. Highlighted points have met significance based on two sample T-test with FDR<0.05 (n=6). (**D**) Proximity ligation assay between DOT1L and chromatin (FLAG tagged histones) in response to 6 h 10 nM romidepsin treatment. Violin plots (left) show quantification of PLA foci per nucleus across 10 images. (**E**) Proximity ligation assay investigating interactions of TRIM33 and chromatin in response to 6 h 10 nM romidepsin treatment. Violin plots (left) show quantification of PLA foci per nucleus across 10 images. (**F**) Western blot of changes in pc293T protein expression over time in response to 10 nM romidepsin. (**G**) Western blot showing changes in histone acetylation over time in response to 10 nM romidepsin (**H**) Western blot of histone H3K79 methylation state overtime in response to 10 nM romidepsin treatment. (**I**) Western blot of PRC1 ubiquitination substrate, H2AK119 overtime in response to 10 nM romidepsin treatment. (**J**) Volcano plot of proximity proteomics of pc293T cells treated with either 10nM romidepsin or DMSO for 24 h. Highlighted points are proteins of interest which have met significance based on two sample T-test with FDR<0.05 (n=6). (**K**) Violin plots of PLA quantification across 10 images of the indicated proteins interaction with FLAG-tagged histones at 24 h treatment of either 10 nM romidepsin or DMSO. (**L**) Western blot of pc293T cells treated with either DMSO or 10 nM romidepsin for 24 h. Stained for pan-phospho S/T or pan-phospho Y. (**M**) Violin plots of PLA quantification across 10 images of ATF3 interactions with known binders c-JUN, c-FOS, and HDAC1 at 24 h treatment of either 10 nM romidepsin or DMSO. (**N**) Phosphoproteomics and MesoMap of B-WICH complex members in pc293T cells treated with 10 nM romidepsin for 24 h. Log_2_FoldChange plotted on the y-axis. For proximity labeling data (blue), positive values are displaced from chromatin and negative values are recruited. For phosphoproteomics (grey), positive values shoe increase in phosphorylation state. Mesomap n=6. Phosphoproteomics n=5. **** indicates P<0.0001 based on two-tailed unpaired t-test. Scale bars = 10 µm.

First, we ensured romidepsin treatment maintains a phenotypic effect on pc293T cells. Concurrent with prior reports, we saw significant changes in histone acetylation when treating at 10 nM as measured by changes in acetylation at H3K9/14 (**Fig. S2A**) *(12)*. Further, pc293T cells were sensitive to romidepsin, leading to a significant decrease in proliferation after a 72h treatment (**Fig. S2B**).

We next sought to determine the romidepsin-induced changes in chromatin occupancy at short (6 h) and long (24 h) treatment times. Our chemoproteomics protocol revealed 23 significant differentially enriched proteins after 6 h, and 1789 significant differentially enriched proteins after 24 h **(Fig. 2C, 2J, and Table S2-5)**. Global changes in nuclear protein abundance were also quantified as a control to ensure that chromatin occupancy, rather than protein expression, was measured using this technique (**Fig. S2C-D and Table S2-5**).

Analysis of enriched proteins at 6 h revealed proteins recruited and displaced from chromatin were consistent with changes in histone acetylation **(Fig. 2C)**. These interactions were validated through proximity ligation assays (PLA), which required antibodies for the proteins of interest and FLAG for the inserted histone constructs, which together showed robust interaction changes that matched our PL **(Fig. 2D-E)**. Western blotting nuclear extracts after romidepsin treatment over the course of 24 h showed no significant change in the expression of top hits MIDEAS, KMT5C, TRIM33, or DOT1L, suggesting our data is measuring changes in physical interactions between chromatin and chromatin modifying proteins, rather than changes in expression **(Fig. 2F)**.

To correlate changes in chromatin occupancy with function, we next assessed changes in post-translational states associated with proteins where romidepsin treatment is known to regulate chromatin binding. We first blotted for various acetylation sites on histone H3, which would correlate HDAC inhibition with recruitment of bromodomain containing proteins, leading to a subsequent loss of methylated lysine readers. In line with literature reports, increased acetylation at histones H3K9/14, K23, and K27 was observed **(Fig. 2G and Fig. S2A)** *(12, 13)*. A robust change in the methylation state of H3K79 was observed over time, consistent with a loss of DOT1L-histone interactions. H3K79me3 was decreased, H3K79me2 was unchanged, and H3K79me, a mark associated with active transcription, was markedly increased **(Fig. 2H)** *(14)*. Analysis of the data for enriched sets of genes that constitute epigenetic complexes (**Fig. S2E**) showed displacement of PRC1.1, a non-canonical polycomb complex that mediates ubiquitylation of H2A via RING1, independent of PRC2 and H3K27me3 *(15, 16)*. Immunoblotting for H2AK119Ub following romidepsin treatment showed a gradual decrease in this mark over 24 h treatment, and no change to H3K27me3 over same time course, signifying a mechanism where PRC1.1 is directed to chromatin through KDM2 **(Fig. 2I)** *(15, 17)*. Thus, MesoMap broadly captures distinct protein complexes, while still retaining sufficient resolution to distinguish an individual protein’s engagement with chromatin, highlighting the utility of our approach.

Comparing chromatin occupancy of epigenetic remodeling complexes over time showed dynamic changes in chromatin interactions. Recruitment of NuRD, SWI/SNF, and ATAC complex members is observed at 6 h, though all members of both complexes were displaced from chromatin by 24 h (**Fig. S2F**). NuA4, Swr1, B-WICH, and Ino80 were progressively displaced from chromatin over time, with small changes observed at 6 h that grew in magnitude over the next 18 h. These data illustrate the platform’s ability to track changes in chromatin occupancy over time in response to extracellular stimulus. Critically, these data cannot be obtained through proximity labeling of any single histone subunit (**Fig. S3** and **Table S6-10**).

### Transcriptional changes in response to HDACi

After a 24 h treatment of romidepsin, we observed significant changes in the chromatin occupancy of several classes of transcription factors, including AP-1, GATA, and the Yamanaka factors SOX2 and KLF3 **(Fig. 2J)**. Recruitment of AP-1 members ATF3, JUN, and JUNB supports observations previously made via ATAC-seq in HDACi treated patient samples *(18)*. Based on these correlations, we validated the recruitment and displacement of a subset of enriched proteins associated with histone acetylation (displaced: P300, HDAC1) and AP-1 transcription (Recruited: JunB, ATF3) using PLA (**Fig. 2K** and **Fig. S4A-D**). Subsequently, we performed secondary validation of JunB association to chromatin using NanoBit (JunB; **Fig. S4E**) and DNA binding activity by ELISA assays (JUNB, c-JUN, c-FOS, ATF3; **Fig. S4F**).

As many transcription factors are regulated via phosphorylation, we questioned whether transcription factor phosphorylation was promoted by romidepsin treatment leading to recruitment to chromatin *(19, 20)*. Western blotting of treated samples showed an increase in both tyrosine and serine/threonine phosphorylation following 24 h treatment **(Fig. 2L)**. Phosphoproteomics analysis provided a total of 2984 nuclear phosphosites with 1954 of these being significantly regulated in response to 24 h romidepsin treatment (**Fig. S5A Table S11-12**). Several members of the AP-1 transcription factor family were phosphorylated in response to romidepsin treatment, including c-JUN (S243, S73), JUND (S90, S100) and ATF3 (T162) **(Fig. S5A-C)**, suggesting romidepsin-induced phosphorylation is responsible for transcriptional regulation of AP-1 genes through recruitment to chromatin and altered cistomes (**Fig. 2M** and **Fig. S5D-F**) *(21, 22)*.

Additionally, correlation of enriched phosphopeptides with proteins enriched in the MesoMap dataset revealed a displacement of the B-WICH complex from chromatin that was associated with increased phosphorylation of 4 of 6 subunits within the complex **(Fig. 2N** and **Fig S5G)**. This finding agrees with prior reports and suggests that this remodeling complex is regulated by phosphorylation (**Fig. S5H**) *(23, 24)*.

### Deconvolution of the orphaned drug candidate SR-1815

The small molecule SR-1815 was recently discovered using a phenotypic screening platform that identifies agents that regulate the abundance of endogenous proteins expressed from pre-selected disease-associated genes within a relevant cellular context. SR-1815 restored endogenous SynGAP protein levels and corrected synaptic and neuronal hyperexcitability, two of the most consequential phenotypes caused by *Syngap1* haploinsufficiency *(Samowitz et al*., *2025*; co-*submitted)*. Because *SYNGAP1* Developmental and Encephalopathic Epilepsy *(SYNGAP1-DEE)* is caused by reduced SynGAP protein expression in brain cells *(25, 26)*, the SR-1815 scaffold is positioned as a potential candidate for treating this genetic neurodevelopmental disorder. However, because SR-1815 is a previously uncharacterized first-in-class small molecule discovered through phenotypic screening, the molecular mechanisms underlying its effects remain unknown, limiting its clinical potential.

To evaluate the suitability of SR-1815 for the MesoMap platform, we first determined how and to what extent this probe regulates transcription. Indeed, we found that SR-1815 dramatically regulates gene expression in cells. For example, treating cortical neurons for 4 h led to hundreds of differentially expressed genes (DEGs), while thousands of DEGs were detected after a two-week treatment **(Table S13-16)**. A workflow that combined RNA-*seq* with standard proteomics revealed that differentially expressed transcripts were highly correlated with accompanying changes in protein abundance **(Figure 3A-B, Fig. S6A**, and **Table S13-17)**. Thus, SR-1815 reorganizes the neuronal proteome by altering transcriptional dynamics. An analysis of biological pathways suggested that both acute and longer-term SR-1815 treatments regulate neuronal development, cytoskeletal dynamics, synapse biology, and mRNA processing, as well as signaling pathways traditionally linked to these cellular processes **(Fig. 3C-E** and **Fig. S6B)**. We confirmed that SR-1815 regulated these cellular processes and neuronal structures. For example, a SynGO analysis indicated that postsynaptic biology was disproportionately impacted by both the short and longer-term SR-1815 treatment **(Fig. 3C-E** and **Fig. S6B)**. Indeed, expression of some postsynaptic proteins, including SynGAP, PSD95, and GluN1 were elevated, while others were not, including several presynaptic proteins **(Fig. 3F-G)**. Consistent with an increase in postsynaptic protein abundance, dendritic spine size was also increased **(Fig. 3H)**. Furthermore, SR-1815 regulated cytoskeletal dynamics linked to neuronal differentiation, as evidenced by more elaborated dendrites with increased total MAP2 signal, a protein expressed within dendritic microtubules **(Fig. 3I-J)**.

**Fig. 3.**
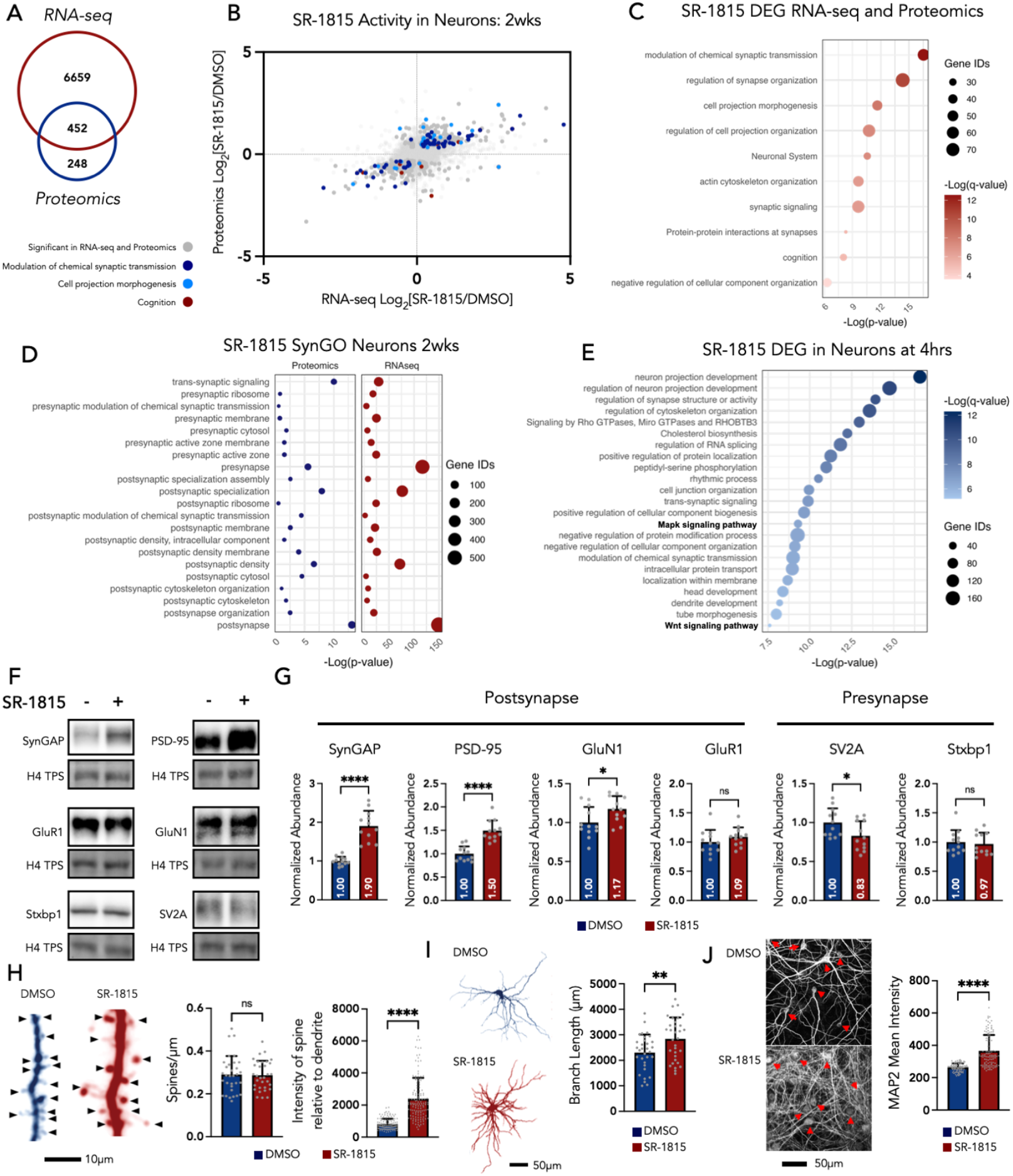
SR-1815 regulates synapse biology through induction of transcriptional dynamics. **(A)** Venn diagram showing the overlap of genes and proteins that passed significance criteria in RNA-seq and proteomics from sister neuronal cultures each treated with SR-1815 for 2 w. Proteomics significance determined by two sample T-test with FDR<0.05 (n=6). RNA-seq significance determined by two sample t-test with FDR <0.01 **(B)** Scatterplot of Log_2_(SR-1815/DMSO) of proteins/genes identified in both RNA-seq (x-axis) and proteomics (y-axis). Highlighted genes are those found in select GO terms from fig. 3C. **(C)** Dotplot of gene ontology of significant differentially expressed proteins identified in both RNA-seq and Proteomics data sets. GO conducted in metascape. Color indicates -Log(q-value) and size indicates number of proteins identified for each GO term. **(D)** Dotplot of presynapse, trans-synaptic and postsynapse related GO terms. Proteomics data (left/blue) and RNAseq data (right/red) plotted separately. -Log(p-value) plotted on x-axis and size indicates number of proteins/genes identified in each GO category. **(E)** Dotplot of gene ontology of differentially expressed genes in mouse neurons treated with SR-1815 for 4 h. Color indicates -Log(q-value) and size indicates number of genes identified for each GO term. Analysis conducted in Metascape. **(F)** Western blots of pre and postsynaptic proteins taken from DIV 14 *Syngap1*^*+/ls*^ neurons treated at plating with vehicle (DMSO) or SR-1815. TPS = Total Protein Stain. **(G)** Quantification of relative intensity of bands normalized to total protein signal. SynGAP: Mann Whitney Test, U=0, P<0.0001, PSD-95: Mann Whitney Test, U=2, P=<0.0001, GluN1: Mann Whitney Test, U=35, P=<0.0332, GluR1: Mann Whitney Test, U=51, P=<0.2415, SV2A: Mann Whitney Test, U=35, P=0.0332, Stxbp1: Mann Whitney Test, U=63, P=0.6297, n=12 per treatment. **(H)** Left: Representative images of spines from eGFP-expressing *Syngap1*^*+/-*^ neurons treated with DMSO or SR-1815 at 1.5 µM for 14 d. Scale bar = 10 µm. Middle: Mean number of spines per dendrite. n=32 per treatment. Mann Whitney test, U=503, P=0.9070. Right: Relative intensity of spine size. n=129 for DMSO, n=130 for SR-1815 Mann Whitney test, U=1080, P=<0.0001. **(I)** Left: Representative dendritic traces of eGFP-expressing *Syngap1*^*+/-*^ neurons treated with DMSO or SR-1815 at 1.5 µM for 14 d. Right: Graph represents quantification of average branch length per neuron. n=32 neurons per treatment. Unpaired t-test P=0.0073 Scale bar=50 µm. **(J)** Left: Representative images showing MAP2 labeling of *Syngap1*^*+/-*^ neurons treated with DMSO or SR-1815 at 1.5 µM for 14 d. Right: Quantification of image mean intensity. N=108 per treatment. Scale bar = 50 µm. Error bars represent standard deviation.

We hypothesized that the transcriptional activity induced by this ligand is mediated, at least in part, though molecular processes that regulate chromatin dynamics. To test this, we performed MesoMap on pc293T cells treated with SR-1815. This revealed significant changes in chromatin dynamics in response to compound treatment **(Fig. 4A** and **Table S18-19)**. For example, 407 proteins were detected in the initial MesoMap analysis, which reflected both increases and decreases in the level of proteins associated with chromatin. However, analysis of the whole proteome under the same conditions revealed changes in only 19 proteins, demonstrating the value of studying mesoscale protein movement as opposed to raw changes in abundance **(Fig. 4B** and **Table S19)**. We observed the recruitment of CDC-like kinases, CLK1/3/4, to chromatin along with RNA splicing machinery, while CLK2 was displaced from chromatin (**Fig. 4A-B**). Moreover, the recruitment of c-Jun and RhoGTPase CDC42 was also observed along with displacement of the calcium regulated kinase CaMKII. These protiens are major regulators of neuronal development and synapse biology and are thus consistent with the observed effects of SR-1815 on dendritic morphogenesis, dendric spine structure, and expression of synapse-targeted proteins *(27– 30)*.

**Fig. 4.**
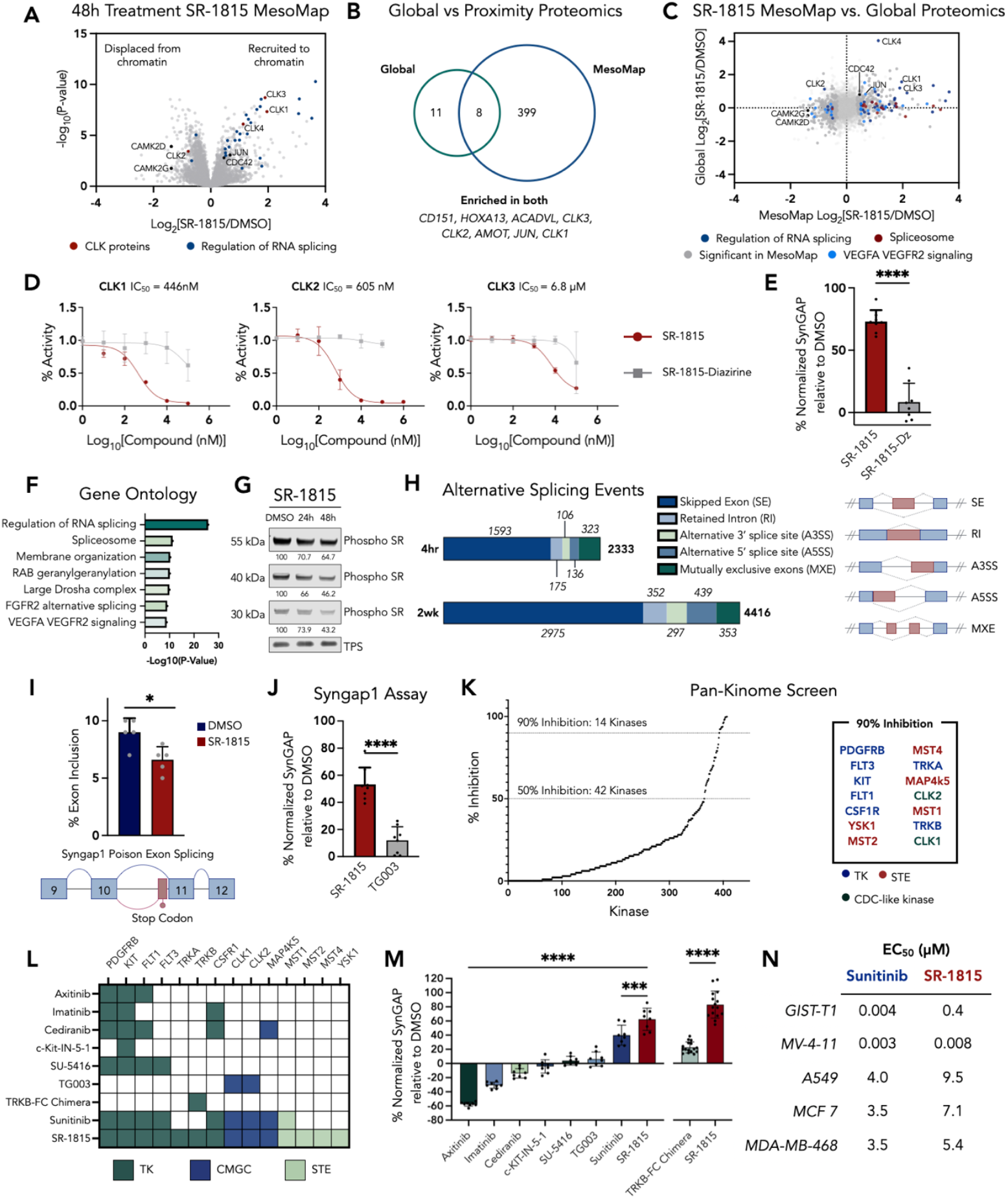
SR-1815 regulates SynGAP expression through kinase inhibition. (**A**) Volcano plot of MesoMap of pc293T cells treated with 10 µM SR-1815 for 48 h. Positive Log_2_FoldChange indicates recruitment to chromatin. CLK proteins highlighted in red and RNA splicing regulators highlighted in blue. Highlighted points have met significance based on two sample T-test with FDR<0.05 (n=6). (**B**) Venn diagram of 407 proteins differentially expressed in global proteomics (FDR < 0.05, n=6) and 19 proteins differentially localized to chromatin in MesoMap (FDR<0.05, n=6). (**C**) Scatterplot of Global (y-axis) and MesoMap (x-axis) proteomics. All points highlighted met significance (FDR<0.05) in MesoMap experiment. Colored points are genes identified with the corresponding GO term. (**D**) In vitro kinase activity showing inhibition of CLK1/2/3 by SR-1815 but not by inactive diazirine-alkyne conjugated SR-1815. Points represent mean and bars represent standard deviation. The connecting line and corresponding IC50 is from a least squares nonlinear regression (n=3). (**E**) SynGAP protein expression level of *Rosa26*^+/fLuc^; *Syngap1*^hb/fl^ neurons treated with SR-1815 or SR-1815-DZ measured using Syngap1 assay, n=8 per treatment. (**F**) Gene ontology of proteins significantly recruited to chromatin in 48 h SR-1815 MesoMap experiment. (**G**) Western blot staining for pan-phospho SR proteins in pc293T cells treated with 10 µM SR-1815 for either 24 or 48 h. Numbers under each blot are percent signal intensity for each band normalized to DMSO. (**H**) Bar graph of significant alternative splicing events occurring in neurons treated with SR-1815 for the indicated duration. Different splicing events are color coded. Illustration of splicing events (right). Differential splicing analysis by rMATS-turbo with FDR<0.05 (n=5). (**I**) Bar graph of *SynGAP1* poison exon inclusion in neurons treated for 2 weeks with SR-1815. Graphing mean and standard deviation of percent spliced in (PSI) of poison exon in each replicate (n=5). (**J**) SynGAP protein expression level of *Rosa26*^+/fLuc^; *Syngap1*^hb/fl^ neurons treated with SR-1815 or TG003 measured using Syngap1 assay, n=8 per treatment. (**K**) Waterfall plot of kinome assay with kinases plotted on x-axis and ordered by % inhibition relative to DMSO (y-axis). Kinases with >90% inhibition are highlighted (right) and colored by kinase family. (**L**) Table showing kinase targets of different known kinase inhibitors tested in neurons for ability to up regulate SynGAP expression. (**M**) SynGAP protein expression level of *Rosa26*^+/fLuc^; *Syngap1*^hb/fl^ neurons treated with kinase inhibitors (n=8) or SR-1815 (n=8), and TRKB-FC Chimera (n=16) or SR-1815 (n=16) measured using Syngap1 assay. (**N**) EC_50_ of Sunitinib and SR-1815 tested in five different cancer cells lines. For panels E, I, J and K **** indicates P < 0.0001, *** indicates P = 0.0009, and * indicates P = 0.0125 based on two-tailed unpaired t-test. For panel M **** indicates P < 0.0001 and *** indicates P = 0.0001 based on one-way ANOVA. Error bars represent standard deviation.

To gain additional insight into how SR-1815 regulates protein dynamics at chromatin, we next overlayed the MesoMap mobility dataset onto the traditional proteomics dataset. This analysis indicated that SR-1815 treatment displaced CLK2 from chromatin while increasing its expression **(Fig. 4C)**, which suggested that SR-1815 may directly regulate CLK2 kinase activity. This was confirmed through a series of NanoGlo kinase assays, where SR-1815 directly inhibited CLK1 and CLK2 activity [IC50 = 446 nM, 605 nM], with significantly less inhibition of CLK3 [IC50= 6.8 µM] (**Fig. 4D**). A diazirine-bearing derivative of SR-1815 that was inactive at CLKs was also inactive in the *Syngap1* expression assay **(Fig. 4E)**.

A principal function of CLKs is to control gene expression dynamics through regulation of splicing *(31)*. These kinases phosphorylate SR-containing proteins, a signal that regulates binding of splice-regulating proteins at chromatin *(32, 33)*. CLK dysregulation is linked to both cancer and neurological disorders, and consequently, CLK inhibitors are being developed for various clinical applications *(31, 33)*. Based on this, we hypothesized that SR-1815 regulates SynGAP expression, at least in part, by altering neuronal splicing. Consistent with this idea, GO terms for genes that exhibited chromatin mobility in response to SR-1815 treatment were enriched for functions related to mRNA processing and splicing **(Fig. 4F)**. SR-1815 caused a decrease in SR protein phosphorylation, consistent with direct CLK1/2 inhibition **(Fig. 4G)**. Critically, the compound dramatically altered neuronal splicing, with a 4 h treatment causing ∼2000 significant splicing events, while more than ∼4000 significant splicing events occurred after 2 w **(Fig. 4H** and **Table S20-21)**. The 4 h SR-1815 treatment induced splicing mainly in genes associated with mRNA processing, while the longer-term treatment induced splicing in genes associated with neuronal morphogenesis and synapse biology (**Fig. S6C**). The time-dependent effect of SR-1815 on neuronal splicing was mechanistically linked to increased SynGAP expression. Only the two-week SR-1815 treatment drove a reduction in the inclusion of an A3SS “poison exon” in *Syngap1* transcripts **(Fig. 4I)** *(34)*. Exclusion of this exon is a mechanism that increases SynGAP protein expression through suppression of nonsense mediated decay of transcript *(34)*. Consistent with this, only the two-week SR-1815 treatment significantly increased *Syngap1* transcript [Log_2_(FC) = 0.18, p-value = 3.52×10^−15^] (**Table S14**). Thus, SR-1815 inhibition of CLK activity regulates SynGAP expression, at least in part, through effects on neuronal splicing.

### SR-1815 is a Multi-Kinase Inhibitor

The findings thus far suggest a mechanism where SR-1815 inhibits CLK1/2, resulting in altered phosphorylation of splicing regulatory complexes, followed by an increase in *Syngap1* levels via reduced inclusion of a *Syngap1* poison exon. Consistent with this, TG003, a selective CLK1/2 inhibitor, enhanced SynGAP expression, but was significantly less efficacious than SR-1815 **(Fig. 4J)** *(35)*. Kinase inhibitors are rarely selective for a single target, suggesting that SR-1815 engages additional kinases. To explore this, SR-1815 was screened against the whole kinome *(36)*. A measure of kinase inhibitor selectivity revealed that this compound exhibits activity across <8% of the kinome (**Fig. 4K, Fig. S7A** and **Table S22)**. This target profile positions it as a relatively selective small molecule kinase inhibitor with similarities to several FDA-approved cancer therapeutics (**Fig. S7B**). As expected, SR-1815 exhibited strong activity at CLK1/2 in the kinome screen **(Fig. 4K** and **Fig. S7A)**. Activity was also observed across 6 kinase families (**Fig. S7B**), with the highest affinity targets belonging to the receptor tyrosine kinase (RTK) family, including KIT, FLT1, FLT3, TRKA/B, PDGRF-b, and CSFR1. In cell kinase occupancy profiling further confirmed SR-1815 binding of kinases (**Fig. S7C-F and Table S23-24**). We mined several databases to identify small molecules known to inhibit one or more targets of SR-1815 and tested their ability to upregulate SynGAP **(Fig. 4L-M)**. Surprisingly, sunitinib exhibited strong activity in the SynGAP expression assay, while the others either had no effect or reduced assay activity **(Fig. 4M)** *(37)*. While these other inhibitors shared at least one common high-affinity target with SR-1815, only sunitinib shared high-affinity targets across multiple kinase families, including RTK, Sterile 20 (STE), and CDKs/ MAPKs/ GSKs/ and CLK (CMGC) **(Fig. 4L** and **Fig. S7B)** *(38)*. This may explain the enhanced efficacy of sunitinib in the SynGAP expression compared to other, more selective, inhibitors. SR-1815 was more efficacious than sunitinib in the SynGAP expression assay (**Fig. 4M**). While SR-1815 is much more selective than sunitinib (**Fig. S7B**), it has unique kinase targets, such as TRKA/B. Indeed, the selective inhibitor of TRKB, TRKB-FC, increased the SynGAP assay signal by ∼20% **(Fig. 4M)**. Thus, the unique targets of SR-1815 likely explain the increased efficacy of SR-1815 relative to sunitinib in the SynGAP expression assay. Finally, the target profile for SR-1815 (**Fig. 4K**), including high-affinity inhibition of both *wildtype* and mutant kinases (**Fig. S7A-B**), suggested that it may demonstrate potent antiproliferative effects in cellular models of cancer. Indeed, it exhibited potent anti-proliferative effects in cancer cell lines that were specifically selected based on a match with SR-1815 high-affinity targets (**Fig. 4N** and **S8A-E)** *(39, 40)*.

## Discussion

Proximity labeling has become a mainstay method in molecular biology for understanding the interactomes of specific proteins in a native environment. Our approach provides evidence that PL can be expanded to intact cellular systems for unbiased discovery of meaningful changes to protein-protein interactions, such as ligand-induced changes to proteins bound to chromatin. This application of PL assigns a molecular fingerprint to a specific cellular stimulus that, in many cases, is more data rich than global analysis of expressed genes. Traditional methodologies used to probe chromatin biology (Co-IP, ChIP-seq, mass-spectrometry, etc.) cannot simultaneously capture change in function of diverse protein complexes throughout chromatin in a single experiment. Beyond proving the approach is possible, MesoMap provided unique molecular insights into how chromatin is regulated in response to HDAC inhibition and predicted the mode of action of the orphan ligand, SR-1815, as a kinase inhibitor that regulates RNA splicing.

The discovery that sunitinib upregulates SynGAP expression supports our conclusion that principal mode of action of SR-1815 is through kinase inhibition. Reciprocal activity displayed by SR-1815 and sunitinib in cancer cell lines and neurons reinforces the biological links connecting tumor biology and neurological disorders, prompting further drug repurposing efforts between both fields. Together, these findings demonstrate how phenotypic screening can be paired with technologies like MesoMap to deconvolute the molecular targets of promising ligands discovered through phenotypic screening, which enables further discovery and development of first-in-class therapeutics for a broad range of diseases.

Future work will focus on adapting this platform into more complex biological contexts to enable universal MesoMap deployment. We expect that extending proximity labeling from single proteins to mesoscale molecular systems will promote novel approaches to studying cellular responses to stimuli and aid in mechanism of action studies for chromatin regulating ligands.

## Acknowledgments

Research reported in this publication was supported by the Office of The Director, of the National Institutes of Health under Award Number S10OD036363, the National Institute of General Medical Sciences of the National Institutes of Health (R35GM150765, R35GM134897) and the National Institute of Mental Health of the National Institutes of Health (R01MH113648, U01MH136567). The content is solely the responsibility of the authors and does not necessarily represent the official views of the National Institutes of Health. CPS also acknowledges the Wertheim UF-Scripps for start-up funds. We would like to thank George Tsaprilis and Catherina Scharager Tapia for assistance with Mass Spectrometry, Robert M. Witwici and Li Pan for assistance with sequencing, and Gogce C. Cryen and Alexander Trouern-Trend for assistance with bioinformatics.

## Author contributions

Conceptualization: CPS, GR, CJD, PS. Manuscript Preparation: CPS, GR, CJD. Experimental Work: CJD, PS, CPS, FT, AL, CB, LR. Data Analysis: CJD, PS, CPS. Funding acquisition and supervision: GR, CPS, DWCM, CM,

## Data and materials availability

All data and materials used in the analysis are available in supplementary materials. Raw mass spectrometry files and RNA sequencing data will be deposited on appropriate publicly available database (i.e. MassIVE and GEO). Plasmids used will be made available via Addgene. Other materials can be made available via request.

