## Supplementary Figures and data for "Mesoscale proximity labeling to study macro changes to chromatin occupancy"

**The PDF file includes:**

Materials and Methods
Figs. S1 to S8
References

### Materials and Methods

#### Materials

Antibodies Used in This Study

| Antibody | Source | Identifier | Use |
|---|---|---|---|
| Goat Anti Rabbit IgG (H+L) Alexa 488 | Invitrogen | A32731 | Western Blot and Immunocytochemistry |
| Goat Anti Rabbit IgG Alexa 555 | Abcam | AB150078 | Western Blot and Immunocytochemistry |
| Goat Anti Rabbit IgG (H+L) Alexa 647 | Invitrogen | A32733 | Western Blot and Immunocytochemistry |
| Goat Anti Rabbit 680 | Li-Cor | 926-68071 | Western Blot |
| Goat Anti Rabbit 800 | Li-Cor | 926-32211 | Western Blot |
| Goat Anti Mouse IgG (H+L) Alexa 488 | Invitrogen | A32723 | Western Blot and Immunocytochemistry |
| Goat Anti Mouse 680 | Li-Cor | 926-68070 | Western Blot and Immunocytochemistry |
| Goat Anti Mouse IgG (H+L) Alexa 647 | Invitrogen | A32728 | Western Blot and Immunocytochemistry |
| Goat Anti Mouse IgG Alexa 555 | Abcam | AB150114 | Western Blot and Immunocytochemistry |
| Streptavidin IRDye 800CW | Li-Cor | 925-32230 | Western Blot |
| HRP-Conjugated Streptavidin | Thermo Scientific | N100 | Western Blot |
| Anti-mouse IgG HRP Conjugate | Promega | W402B | Western Blot |
| Anti-rabbit IgG HRP Conjugate | Promega | W401B | Western Blot |
| Revert 700 Total Protein Stain | Li-Cor | 926-11021 | Western Blot |
| Revert 520 Total Protein Stain | Li-Cor | 926-10021 | Western Blot |
| JunB Mouse monoclonal mAb | Santa Cruz Biotechnology | C-11, sc-8051 | Western Blot and Immunocytochemistry |
| DYKDDDDK (Flag) 9A3 Mouse mAb | Cell Signaling Technology | 8146S | Western Blot and Immunocytochemistry |
| HA Tag Rabbit mAb (C29F4) | Cell Signaling Technology | 3724S | Western Blot and Immunocytochemistry |
| Histone H2BK120 Ubiquitin 1 | Active Motif | 39624 | Western Blot |
| Histone H3K18Ac Mark | Active Motif | 39756 | Western Blot |
| Histone H3K4 Trimethyl Mark | Active Motif | 39160 | Western Blot |
| Di-Methyl-Histone H3 (K4) (C64G9) Rabbit mAb | Cell Signaling Technology | 9725T | Western Blot |

| | | | |
|---|---|---|---|
| Di-Methyl-Histone H3 (K27) (C36B11) Rabbit mAb | Cell Signaling Technology | 9733T | Western Blot |
| Tri-Methyl-Histone H3 Lys9 (D4W1U) Rabbit mAb | cell Signaling Technology | 13969T | Western Blot |
| Histone H3 (D1H2) XP(R) Rabbit mAb | Cell Signaling Technology | 4499S | Western Blot |
| Histone H3 (D1H2) XP(R) Rabbit mAb | Cell Signaling Technology | 4499S | Western Blot |
| Acetyl-Histone-H3 (K9/K14) Rabbit mAb | Cell Signaling Technology | 9677S | Western Blot |
| HDAC1 (D5C6U) XP(R) rabbit mAb | Cell Signaling Technology | 34589T | Western Blot and Immunocytochemistry |
| HDAC1 (10E2) Mouse mAb | Cell Signaling Technology | 5356T | Western Blot and Immunocytochemistry |
| Tri-mETHYL-Histone H3 (K36) (D5A7) XP(R) Rabbit mAb | Cell Signaling Technology | 4909T | Western Blot |
| Di-Methyl-Histone H3 (K9) (D85B4) XP(R) Rabbit mAb | Cell Signaling Technology | 4658T | Western Blot |
| Ubiquityl Histone H2A K119 D27C4 | Cell Signaling Technology | 8240S | Western Blot |
| P-HISTONE H2AX S139 Y142 Rabbit | Cell Signaling Technology | 5438S | Western Blot |
| C-JUN mAb (5B1) | invitrogen | MA5-15881 | Western Blot and Immunocytochemistry |
| JUND pAb | Active Motif | 61403 | Western Blot |
| JUNB (C37F9) Rabbit mAb | Cell Signaling Technology | 3753S | Western Blot and Immunocytochemistry |
| c-JUN (60A8) Rabbit mAb | Cell Signaling Technology | 9165S | Western Blot and Immunocytochemistry |
| CDK1 Polyclonal antibody | Proteintech | 10762-1-AP | Western Blot |
| CDK9 (C12F7) Rabbit mAb | Cell Signaling Technology | 2316S | Western Blot |
| CLK1 pAB | Invitrogen | PA5-112388 | Western Blot |
| Anti ELMSAN1 (MIDEAS) | Sigma | HPA003111 | Western Blot |
| ATF3 pAb | Invitrogen | PA5-106898 | Western Blot, ELISA, and Immunocytochemistry |
| C/EBP beta mAb (1H7) | Invitrogen | MA1-827 | Western Blot |
| DOT1L (D1W4Z) n Rabbit | Cell Signaling Technology | 77087S | Western Blot and Immunocytochemistry |
| FOS-B (5G4) Rabbit mAb | Cell Signaling Technology | 2251S | Western Blot |
| c-FOS (F6) Rabbit mAb | Cell Signaling Technology | 2250S | Western Blot and Immunocytochemistry |

| | | | |
|---|---|---|---|
| c-FOS Mouse | Abcam | AB-208942 | Western Blot and Immunocytochemistry |
| SUV420H2 pAb (KMT5C) | Invitrogen | PA5-115484 | Western Blot and Immunocytochemistry |
| P300 pAb | Invitrogen | PA1-848 | Western Blot and Immunocytochemistry |
| (P-S/T) Phe Rabbit Ab | Cell Signaling Technology | 9631S | Western Blot |
| P-Y-1000 Multiman (TM) Rabbit mAb mix | Cell Signaling Technology | 8954S | Western Blot |
| SRPK1 Rabbit | Proteintech | 14073-1-AP | Western Blot |
| TFAP4 pAb | Invitrogen | PA5-51658 | Western Blot |
| TBP (D5C9H) XP(R) Rabbit mAb | Cell Signaling Technology | 44059S | Western Blot |
| TRIM33 d7U4F rabbit mAb | Cell Signaling Technology | 90051S | Western Blot and Immunocytochemistry |
| DYKDDDDK Tag | Epicypher | 13-2031 | CUT&Tag |
| Histone H3K27ac | Active Motif | 39134 | CUT&Tag |
| Histone H3K27me3 | Active Motif | 39157 | CUT&Tag |
| Guinea Pig Anti-Rabbit | Active Motif | 105465 | CUT&Tag |
| IgG Rabbit | Cell Signaling Technology | 27295 | CUT&Tag |
| H3K20ac pAb | Invitrogen | 720087 | Western Blot |
| Histone H3K79me1 pAb | Epigentek | C10009-1 | Western Blot |
| Histone H3K79me2 pAb | Epigentek | C10009-1 | Western Blot |
| Histone H3K79me3 | Epigentek | C10009-1 | Western Blot |
| Histone H4K20 Monomethyl pAb | Epigentek | A-4046-025 | Western Blot |
| Histone H4K20 Dimethyl pAb | Epigentek | A-4047-025 | Western Blot |
| Histone H4K20 Trimethyl pAb | Epigentek | A-4048-025 | Western Blot |
| Histone H3K4me1 pAb | Epigentek | C10005-1 | Western Blot |
| Histone H3K4me2 pAb | Epigentek | C10005-1 | Western Blot |
| Histone H3K4me3 pAb | Epigentek | C10005-1 | Western Blot |
| Anti-acetyl-Histone H3 (Lys23) | Invitrogen | 07-355 | Western Blot |
| Pan-SynGAP antibody | Cell Signaling Technology | 5539 | Western Blot |
| PSD-95 antibody | Addgene | 180082-rAb | Western Blot |
| GluN1 antibody | Addgene | 182200-rAb | Western Blot |
| GluR1 antibody | Addgene | 180093-rAb | Western Blot |

| | | | |
|---|---|---|---|
| SV2A antibody | Cell Signaling Technology | 66724 | Western Blot |
| Stxbp1 (Munc18-1) | Abcam | AB124920 | Western Blot |

Compounds Used

| **Compound** | **Manufacturer** | **Catalog #** |
|---|---|---|
| Imatinib | Cayman Chemical | 13139 |
| Cediranib | Cayman Chemical | 11495 |
| TG003 | Cayman Chemical | 10398 |
| Sunitinib | Cayman Chemical | 13159 |
| TRKB-FC Chimera | biotechne | 688-TK-100 |
| SU 5416 | biotechne | 3037/10 |
| Sunitinib malate | biotechne | 3768/10 |
| Axitinib | biotechne | 4350/10 |
| Cediranib | biotechne | 7454/10 |
| KN-62 (CAS 127191-97-3) | Santa Cruz | sc-3560 |
| c-Kit-IN-5-1 | Medchem express | HY-18302 |
| TG003 | Medchem express | HY-15338 |
| Pexidartinib | Medchem express | HY-16749 |
| Romidepsin | Sigma Aldrich | SML1175 |

**Methods**

Cell Culture

HEK 293T cells were cultured as a monolayer in DMEM (Gibco #11995073), supplemented with 10% v/v FBS (R&D Systems S11150), 100 U $mL^{-1}$ of penicillin 100 µg $mL^{-1}$ of streptomycin (Gibco 15140-122). Cells were maintained in an incubator at 37 °C with 5% CO2. Pc293T cells were grown in complete DMEM media (above) with the addition of 400 µg/mL Zeocin (Invitrogen 46-0072) and 100 µg/mL Hygromycin (Invitrogen 10687010).

Stable cell line generation

The pACEMAM3 Integrator Module (Geneva Biotech) bearing H2A-HA-CfaN-FLAG, H3.1-HA-CfaN-FLAG, H3.3-HA-CfaN-FLAG, and H4-HA-CfaN-FLAG was linearized using PI-PspI restriction enzyme (New England Biolabs R0695S). DNA was electrophoretically separated through a 1% agarose gel, linearized bands were excised, and DNA was purified using the Monarch DNA gel extraction kit (New England Biolabs T1020L). 3.5 µg of linearized plasmid was transfected into 60% confluent HEK293T cells grown in a 10 cm plate using Lipofectamine 2000 (ThermoFisher Scientific 11668027) and incubated for 24 h. Cells underwent selection with media containing 100 µg/mL hygromycin (Invitrogen 10687010) and 400 µg/mL Zeocin (Gibco R25001). After recovery, cells underwent FACS performed by the flow cytometry core at The Herbert Wertheim UF Scripps Institute for Biomedical Innovation and Technology on a BD FACSAria Fusion (BD biosciences). Single cells expressing GFP in top 10% of cell population were plated in single cells per well in a 96-well plate. Colonies were expanded while continuing antibiotic treatment.

Flow Cytometry

$1x10^6$ cells were collected with trypsin and quenched with complete media. Cells were pelleted at 400xg for 3 min, supernatant was removed, and cells were washed with DPBS. Cells were pelleted and resuspended in sort buffer (2.5 mM EDTA, 25 mM HEPES pH 7.0, 1% FBS) and passed through 35 µm strainer. Cells were analyzed using a BD LSR II (BD Biosciences) and gated for live cells and singlets using forward and side scatter characteristics and GFP expression was captured using 488 nm laser. A minimum of 10,000 events were captured per sample. GFP signal was gated using wild type HEK293T cells as negative control. FlowJo software was used to analyze data and create figures.

Transient Transfection

For single histone construct experiments HEK293T cells grown to 60% confluence in 10 cm plates were transfected with 3.5 µg of plasmid DNA using Lipofectamine 2000 and incubated overnight. Media was changed to complete media and cells were treated with 10 nM romidepsin or DMSO for 24 h and collected.

Western Blot

Cell lysates were quantified using Pierce BCA protein assay kit (Thermo Scientific #A55864) and normalized to same concentration. Samples were electrophoretically separated with a pre-cast Invitrogen NuPAGE 4-12% bis-tris acrylamide gel (Invitrogen #NW04125BOX) and transferred to nitrocellulose membrane. Samples were then stained with Revert 520 total protein stain (LICOR Bio 926-10011) and imaged on a Licor Odyssey M. The membranes were then blocked with 2% BSA in TBST for 1 h at room temperature and stained with primary antibody diluted in TBST overnight at 4 °C (dilutions based on antibody manufacturer recommendations). Membranes were washed 3x with TBST and then stained with secondary fluorescent antibody diluted 1:10,000 in TBST for 1 h at room temperature (Licor: 926-68070 and 926-32211; Invitrogen: A32787 and A32731). Membranes were then washed 3x with TBST and imaged on Licor Odyssey M. Quantification of western blots was performed in Fiji.

Immunocytochemistry

Cells were cultured, plated, fixed and permeabilized as described for proximity ligation assays. Coverslips were then blocked with 1% BSA in PBS for 1 h at room temperature. Coverslips were then incubated with primary antibody diluted 1:200 in 1% BSA in PBS overnight at 4 °C. Samples were then washed 3x with PBS and incubated with secondary antibody diluted 1:1000 for 1 h at room temperature. Slides were then washed 3x with PBS and mounted to slides using Vectashield DAPI mounting media and sealed with clear nail polish. Slides were imaged as described for the proximity ligation assays.

CUT&TAG

CUT&Tag was performed using the CUT&Tag-IT® Assay Kit (Active Motif # 53160). The manufacturer protocol was followed with the application of the following antibodies: Histone H3K27me3 (Active Motif #39157), H3K27ac (Active Motif #39134), DYKDDDDK Tag (EpiCypher #13-2031), and Rabbit IgG (Cell Signaling #2729S). Each target was run in triplicate with 2x10$^5$ pcHEK293T cells per sample. Sequencing was conducted on a NextSeq 2000 at 35m reads per sample by The Herbert Wertheim UF Scripps Institute for Biomedical Innovation & Technology Genomics Core.

CUT&TAG data analysis

Samples were processed using nf-core/cutandrun (https://nf-co.re/cutandrun/3.2.2/). Samples were aligned to GRCh38.p14 and the following changes were made to the nf-core pipeline: -profile singularity --peakcaller macs2 --remove_mitochondrial_reads --skip_trimming --skip_removeduplicates --normalisation_mode CPM. Tracks were viewed in IGV, and heatmaps were generated using deepTools.

Proximity Labeling

For cells treated with compound, all lysis buffers contained compound or vehicle at same concentration as treatment. A total of $3 \times 10^7$ cells expressing POI-HA-CfaN-FLAG were resuspended in 1 mL PBS and then lysed with addition of 4 volumes of ice-cold lysis buffer (45 mM KCl, 0.1 mM EDTA, 6.25 mM $MgCl_2$, 12.5 mM Tris, 375 mM Sucrose, 0.125% Nonidep P40, 1x Halt Protease Inhibitor Cocktail EDTA-free Thermo Scientific #1861279) for 10 min on ice. The crude nuclei were isolated by centrifugation at 400xg for 5 min at 4 °C. The nuclei were washed 3x with DPBS + Halt Protease Inhibitor and centrifugation at 400g for 5 min to pellet nuclei. The nuclei were resuspended in 600 µL of DPBS with 0.25 µM CfaC-Ir or CfaC-biotin. The nuclei were incubated at 37 °C for 30 min with gentle end-over-end rotation.

The nuclei were isolated by centrifugation at 400xg for 5 min at 4 °C and washed twice with DPBS (500 µL) to remove excess peptide. The pellets were then resuspended in 1 mL of DPBS containing diazirine–biotin conjugate (200 µM) and irradiated with blue light for 1 min in the Penn PhD Photoreactor M2 at 100% light intensity at 4 °C. The nuclei were re-isolated by centrifugation at 400xg for 5 min at 4 °C and washed once with DPBS to remove excess biotin–diazirine. The washed pellets were then resuspended in 1 mL of LB3 buffer (10 mM Tris, 100 mM NaCl, 1 mM EDTA, 0.5 mM EGTA, 0.1% sodium deoxycholate, 0.5% sodium lauroyl sarcosinate, pH 7.5) and sonicated using Diagenode Bioruptor Plus for 14 cycles of 30 sec on and 30 sec off at max power and 4 °C.

Streptavidin enrichment

The lysed nuclei were then clarified through centrifugation at 15,000xg for 20 min at 4 °C and the protein concentration of the supernatant was determined by BCA assay. Protein concentration was normalized across all experimental replicates and diluted to 1 mg/mL with LB3. Then 1 mL of each sample was incubated with 125 µL of prewashed magnetic Sepharose streptavidin beads (Cytiva Life Sciences no. 28985738) for 2 h at room temperature or overnight at 4 °C with end-over-end rotation. The beads were subsequently washed 3x with 1% w/v SDS in PBS, 3x with 1 M NaCl in PBS and 3x 10% ethyl alcohol in PBS for 5 min each wash with end-over-end rotation.

Label Free Proximity Proteomics

Following streptavidin enrichment, the beads were resuspended in PBS (500 µL) and transferred to a new 1.5 mL LoBind tube. The supernatant was removed, and the beads were washed with 3× PBS (0.5 mL) and 3× ammonium bicarbonate (100 mM). The beads were resuspended in 500 µL of 3 M urea in PBS and 25 µL of 200 mM dithiothreitol in 25 mM $NH_4HCO_3$ was added. The beads were incubated at 55 °C for 30 min. Subsequently, 30 µL of

500 mM iodoacetamide in 25 mM $NH_4HCO_3$ was added and incubated for 30 min at room temperature in the dark. The supernatant was removed and the beads washed with 3x with 0.5 mL of DPBS and 6x with 0.5 mL of 50 mM ammonium bicarbonate. The beads were resuspended in 0.5 mL of ammonium bicarbonate (50 mM) and transferred to a new protein LoBind tube. The beads were resuspended in 40 µL of ammonium bicarbonate (50 mM), 1.2 µL of trypsin (1 mg/mL in 50 mM acetic acid) was added and the beads incubated overnight with end-over-end rotation at 37 °C. After 16 h, a further 0.8 µL of trypsin was added and the beads were incubated for an extra 1 h at 37 °C. Supernatant was then transferred to new LoBind tubes and each biological replicate split into two technical replicates.

Label Free Nuclear Proteomics

After splicing and lysis of nuclei in LB3, 20 µg of protein was taken for cleanup and digestion. Proteins was cleaned up and digested using Sera-Mag Carboxylate SpeedBeads (Cytiva Life Sciences: E7 Cat. #45152105050250 and E3 Cat. #65152105050250) following manufacturer's protocol. Digested peptides were split into two technical replicates per sample.

Label Free Proteomics Data analysis

Peptide digests were acidified with TFA to 0.1% (v:v) and desalted using 2 µg capacity ZipTips (Millipore, Billerica, MA) according to manufacturer instructions. Following drying under vacuum, peptides were re-solubilized in 20 µL of 0.1% formic acid (FA) to a final concentration of 100 ng/µL. Samples were analyzed on a nanoElute (plug-in V2.1.60.0; Bruker, Germany) coupled to a Bruker TimsTOF Pro 2 mass spectrometer (Breman, Germany), equipped with a CaptiveSpray source and a 20 µm zero dead volume (ZDV) Sprayer. Peptides (corresponding to 400 ng) were separated on a reverse-phase C18 column (10 cm X 75 µm X 1.9 µm, Bruker PepSep Ten Series). The column temperature was maintained at 50 °C using an integrated Bruker Column Toaster (Bremen, Germany). The column was equilibrated using 4 column volumes before loading samples in 100% buffer A (99.9% Fisher Optima® LC/MS water, 0.1% FA), with both steps performed at 800 bar. Samples were separated at 500 nL/min using a linear gradient from 3% to 30% buffer B (99.9% Fisher Optima® LC/MS acetonitrile, 0.1% FA) over 17.90 min before ramping to 95% buffer B (0.5 min) and sustained at 95% buffer B for 2.4 min (total separation method time 20.7 min). The Bruker TimsTOF Pro 2 was operated in DIA-PASEF mode using Tims Control v. 5.0.2. Settings for the MS method were as follows: Mass Range 100 to 1700 m/z, 1/K0 Start 0.6 V·s/cm$^2$ End 1.4 V·s/cm$^2$, TIMS Ramp and accumulation time 75 ms, Capillary Voltage 1700 V, Dry Gas 3 L/min, Dry Temp 200 °C, DIA-PASEF settings: 18 MS/MS scans (50 m/z windows, 0.21 1/K0 windows, total cycle time 0.74), mass range 300 to 1200, and CID collision energy 20 eV (at 0.60, 1/K0) to 65 eV (at 1.60, 1/K0). The analysis was performed at The Herbert Wertheim UF Scripps Institute for Biomedical Innovation & Technology, Mass Spectrometry and Proteomics Core Facility (RRID:SCR_023576).

Data was processed via DIANN 1.8.1. Parameters set as follows: trypsin/P digestion, 3 missed cleavages, 3 max. variable modifications, N-term M excision, Ox(M), Ac(N-term) and C carbamidomethylation. Peptide length range was 7-30, precursor charge range 1-4, m/z range 300-1800, and fragment ion range 200-1800. Mass accuracy and MS accuracy were both set to 10. The following settings on the algorithm were checked: "Use isotopologues", "MBR", "No shared spectra", "Heuristic protein inference". Precursor FDR was set to 1%. A spectral library was used generated via DIANN from all known human proteins (In-Silico spectral library). The resulting matrix.pg file was opened in Perseus (v2.0.7.0). Intensities were inputted as "main", the rest of the descriptors are categorical. Data was then transformed ($Log_2$). Data is annotated by treatment. Missing values were imputed with Perseus default settings. Normalization was performed via median subtraction. Following this process, a volcano plot was generated utilizing a t-test for statistical significance. The resulting volcano plots were plotted in GraphPad Prism 10 for final figures. Metascape was used for Gene Ontology. Panther database was used to identify complex members.

Phospho Proteomics Experimental Protocol

Cells treated with 10 nM romidepsin for 24 h were collected by trypsinization. Nuclear isolation was conducted as above and subsequently lysed with LB3. All buffers used were supplemented with 5 mM sodium butyrate and Halt Protease and Phosphatase Inhibitor Cocktail (100X) (ThermoFisher Scientific #78440). Protein solution samples (2 experimental conditions, 5 biological replicates/condition) were precipitated overnight with 4X ice cold acetone (v:v). Protein pellets were then solubilized in 100 mL of 20% SDS and water was then added to bring the final solution to 400 mL containing 5% SDS. A Pierce BCA assay (Thermo Fisher Scientific, Watlham, MA) was then performed to determine protein amounts in each sample. Protein solutions (anywhere between 358.9 – 440.3 mg total protein) were then processed for trypsin digestion using Midi S-TrapsTM (Protifi, Huntington, NY) according to the manufacturer's instructions. Briefly, 370 mL of protein solution were reduced using tris-2(-carboxyethyl)-phosphine (TCEP) at 55 °C for 15 min, alkylated using methyl methanethiosulfonate (MMTS) at room temperature for 10 min, and then digested with 20 mg trypsin at 47 °C for 1 h. Following this incubation, 500 µL of 50 mM triethylammonium bicarbonate (TEAB) was added to the Midi S-TrapTM and the peptides were eluted using centrifugation. Elution was repeated once. A third elution using 500 mL of 50% acetonitrile (ACN). The eluted peptides were then dried under vacuum. The dried peptides were resuspended in 50 mM TEAB and their concentrations were determined using the PierceTM quantitative fluorometric peptide assay (Thermo Fisher Scientific, Waltham, MA). One hundred micrograms of peptides/sample were labelled with TMT labels (10-plex) according to the manufacturer's instructions (Thermo Fisher Scientific, Waltham, MA) and pooled. Following labeling the pooled sample was dried again under vacuum and subsequently desalted using Waters Oasis™ OASIS HLB 1 cc solid phase extraction cartridges according to the manufacturer's instructions (Waters, Milford, MA). Five hundred micrograms of material after elution (half the volume), were enriched for phosphopeptides using the High- Select™ Fe-NTA Phosphopeptide Enrichment Kit from Thermo Fisher Scientific (Thermo Fisher Scientific, Waltham, MA) according to the manufacturer's instructions.

Phospho Proteomics Data Analysis

Peptide digest samples corresponding to the phoshopeptide enriched proteome and the flow through proteome following the enrichment, were dried under vacuum and subsequently acidified using 50 mL of 1% TFA (pH <3). The samples were then desalted using NuTip Carbon tips from Glygen (Glygen Corp., Columbia, MD) in the case of phospho peptides, or 2 µg capacity ZipTips (Millipore, Billerica, MA), according to manufacturer instructions. They were then dried under vacuum. Peptides resolubilized in 5 mL of 0.1% TFA were on-line eluted into a Fusion Tribrid mass spectrometer (Thermo Scientific, San Jose, CA) from an EASY PepMapTM RSLC C18 column (2 µm, 100 Å, 75 µm x 50 cm, Thermo Scientific, San Jose, CA), using a gradient of 5-25% solvent B (80/20 acetonitrile/water, 0.1% formic acid) in 180 min, followed by 25-44% solvent B in 60 min, 44-80% solvent B in 0.1 min, a 5 min hold of 80% solvent B, a return to 5%

solvent B in 0.1 min, and finally a 10 min hold of 5% solvent B. The gradient was then extended for the purpose of cleaning the column by increasing solvent B to 100% in 3 min, a 100% solvent B hold for 10 min, a return to 5% solvent B in 3 min, a 5% solvent B fold for 3 min, an increase of solvent B to 100% in 3 min, a 100% solvent B hold for 10 min, a return to 5% solvent B in 3 min and a 5% solvent B hold for 3 min and finally, another increase to 100% solvent B in 3 min and a hold of 100% solvent B for 10 min. All flow rates were 250 nL/min delivered using an nEasy-LC1000 nano liquid chromatography system (Thermo Scientific, San Jose, CA). Solvent A consisted of water and 0.1% formic acid. Ions were created at 1.9 kV using an EASY Spray source (Thermo Scientific, San Jose, CA) held at 50 °C. A synchronous precursor selection (SPS)-MS3 mass spectrometry method was selected based on the work of Ting et al. (Ting, L.; Rad, R.; Gygi, S.P.; Haas, W. MS3 eliminates ratio distortion in isobaric multiplexed quantitative proteomics. Nature Methods 2011, 8, 937–940). Scans were conducted between 380-2000 m/z at a resolution of 120,000 for MS1 in the Orbitrap mass analyzer at an AGC target of 4E5 and a maximum injection of 50 msec. Collision induced dissociation (CID) was then performed in the linear ion trap of peptide monoisotopic ions with charge 2-8 above an intensity threshold of 5E3, using a quadrupole isolation of 0.7 m/z and a CID energy of 35%. The ion trap AGC target was set to 1.0E4 with a maximum injection time of 50 msec. Dynamic exclusion duration was set at 60 sec and ions were excluded after one time within the +/- 10ppm mass tolerance window. The top 10 MS2 ions in the ion trap between 400-1200 m/z were then chosen for Higher-energy C-trap dissociation (HCD) at 65% energy. Detection occurred in the Orbitrap at a resolution of 60,000 and an AGC target of 1E5 and an injection time of 120 msec (MS3). All scan events occurred within a 3 sec. specified cycle time. The analysis was performed at The Herbert Wertheim UF Scripps Institute for Biomedical Innovation & Technology, Mass Spectrometry and Proteomics Core Facility (RRID:SCR_023576).

Phospho(STY) sites file was loaded into Perseus. Rows were filtered based on "Reverse", "Potential contaminant", and localization probability > 0.75. Data was then transformed ($Log_2$). Data table was lengthened using "Expand site table". Samples were annotated by treatment using categorical annotation. Rows were filtered by rows containing > 50% valid values in each group. Imputation was performed using Perseus default settings. Samples were normalized by median subtraction. A volcano plot was generated utilizing a t-test for statistical significance. The resulting volcano plots were plotted in GraphPad Prism 10 for final figures. Kinase activity was predicted using RoKAI App (https://rokai.io/). Spectrum were visualized using PDV (https://github.com/wenbostar/PDV?tab=readme-ov-file).

Proximity Ligation Assay

Round coverslips were place in a 12-well plate and washed with 100% ethanol. Wells were then washed with $H_2O$ and treated with polylysine (Sigma Aldrich #P4707) for 10 min, followed by two washes with $H_2O$ and allowed to airdry for 2 h. Cells were then plated at $2x10^5$ cells/well and allowed to adhere overnight. Cells were then treated with 10 nM romidepsin or vehicle and incubated for desired time. At the indicated time, the media was aspirated, the wells were washed 2x with DPBS and fixed for 10 min with 4% Formaldehyde in DPBS at room temp. Cells were then washed 2x with DPBS, and permeabilized with pre-chilled 100% MeOH for 5min at -20 °C. Slides were washed 3x with DPBS. Once fixed and permeabilized proximity ligation was carried out as recommended by manufacturer using the Duolink in situ red starter kit mouse/rabbit (Sigma-Aldrich DUO92101). In brief, slides were blocked for 1 h at 37 °C with blocking buffer. Slides were then incubated with primary antibodies diluted 1:200 in antibody binding buffer at 4 °C overnight. Slides were washed with wash buffer A and then incubated with PLA probe plus donkey anti-rabbit IgG and PLA probe minus donkey anti-mouse IgG antibodies for 1 h at 37 °C. Samples were washed with wash buffer A and incubated for 30 min at 37 °C with Duolink ligase reaction solution. Slides were washed with wash buffer A and incubated for 100 min with duolink amplification solution. Slides were washed with wash buffer B and mounted with duolink in situ mounting media with DAPI and sealed with clear nail polish.

Slides were imaged on a Nikon Eclipse Ti2 with a CFI60 Plan Apochromat Lambda D 100x Oil Immersion Objective Lens, N.A. 1.45, W.D. 0.13 mm, F.O.V. 25 mm, DIC, Spring Loaded. Cells located using DAPI channel to eliminate bias for PLA signal. Z-stack images were taken with with DAPI, channel 488, and channel 550 using 0.2 µm z-spacing between 11 total images. At least 5 images were taken of each condition and quantified using CellProfiler nuclear speckle counting software. Statistical significance was determined by an unpaired t-test of foci per nuclei using Graphpad Prism with outliers excluded to limit bias from staining inconsistencies.

Transcription Factor Binding Assay

DNA binding activity of AP-1 members was determined using a commercially available TransAM AP-1 DNA-Binding ELISA (Active Motif #44296). The assay was conducted based on manufacturer recommendations. All buffers contained either 10 nM romidepsin or DMSO. To detect ATF3 DNA binding activity an anti-ATF3 antibody was used (Invitrogen PA5-106898). 3 replicates were run per treatment. Absorbance was read 3x at 450 nm and 650 nm. Reads were averaged and 450 nm reads were normalized to the 650 nm background. Samples were then normalized to DMSO = 1. Statistics were calculated with a two-way ANOVA in GraphPad Prism.

NanoBiT Assay

NanoBiT data was run using the NanoBiT® PPI Starter Systems kit (Promega #N2014). H2A and JUNB were cloned into pBiT1.1-N [TK/LgBiT] and pBiT2.1-C [TK/SmBiT] vectors, respectively. HEK293T cells were plated at 10,000 cells/well in a 96-well clear bottom, white walled, tissue culture assay plate (Corning #3610) and cultured overnight. Cells were then transfected with 50 ng of both plasmids per well with lipofectamine 2000 (ThermoFisher Scientific #11668019) in FluoroBrite DMEM and incubated overnight. Cells were then treated with serial dilution of romidepsin or DMSO for 24 h (n=3). To read out luminescence, 10x Nano-Glo live cell substrate was added to each well and incubated for 15 min at 37 °C. Luminescence (signal) and fluorescence (background) were read out on a BioTek Synergy H1 microplate reader 3x for each plate. The average of the reads per sample was taken and the luminescence signal was normalized to background fluorescent signal. Samples were then normalized to vehicle treated samples by division. Results were plotted on Graphpad Prism and statistics were calculated using a One-Way ANOVA.

In Vitro Kinase Assay

The assay of CLK1/2/3 inhibition by SR-1825 was conducted using BPS Bioscience Chemi-Verse Kinase Assay Kits for the respective CLK (#82145) paired with ADP-Glo Kinase Assay (Promega #V6930), following the manufacturer's protocol. Compounds were diluted in a 10x serial dilution from 1 mM to 1 nM and run in triplicate along with a negative control (no recombinant CLK) and a positive control (vehicle). Luminescence was read 3x per plate on a BioTek Synergy H1 microplate reader. Average of reads was normalized to the positive control and fit to a curve using non-linear least squares analysis in Prism Graphpad.

Cell Viability Assay

Cells were plated with 10,000 cells/well in a clear bottom, white walled, 96-well plate and cultured overnight to adhere. The following day, media was removed and replaced with drug at indicated concentration and cultured for 72 h. At end of incubation, media was removed and replaced with complete 1x celltiter-fluor reagent (Promega #G6080) diluted in fluorbrite DMEM (Gibco #A1896701) and incubated for 30 min. Plates were then read 3x for fluorescence on a BioTek Synergy H1 microplate reader. The average of the reads was normalized to vehicle treated wells and fit to a curve using non-linear least squares analysis in Prism Graphpad.

Kinase Occupancy Assay

Kinase occupancy experiments were conducted according to previous reports from Taunton (*41*). In brief, pc293T or neurons were treated with either SR-1815 (10 µM) or vehicle at 37 °C for 3 h. After initial incubation, cells were treated with alkyne probe (2 µM) for 1 h. Cells were washed 1x with warm PBS and lysed in their plates with 100 mM HEPES pH 7.5, 150 mM NaCl, 0.1% NP-40, 1 mM phenylmethyl sulfonyl fluoride (PMSF), 1X complete EDTA-free protease inhibitor cocktail (Thermo Scientific #1861279). Imine reduction was performed by adding 5 mM NaBH4 for 30 min on ice. Lysates were cleared by centrifugation (16,000xg, 30 min, 4 °C). Protein concentration was determined by the BCA assay (Thermo Scientific #A55864). Cell lysates were normalized to 2 mg/mL protein with lysis buffer. Lysates were incubated with 125 µL of streptavidin beads (Cytiva Life Sciences no. 28985738) at 4 °C overnight to remove endogenous biotinylated proteins. The supernatant (mL) was reacted with 191 µL of click chemistry cocktail, resulting in a final concentration of 1% SDS, 100 µM biotin DMTP picolyl azide, 1 mM TCEP, 100 µM TBTA (from a 2 mM stock prepared in 1:4 DMSO:t-butyl alcohol) and 1 mM $CuSO_4$. After incubation at room temperature for 90 min, the proteins were precipitated by adding 10 mL of prechilled acetone and incubating overnight at –20 °C. The precipitated proteins were pelleted by centrifugation (3500g, 4 °C, 30 min), resuspended in cold methanol and re-pelleted. The pellet was solubilized in 1% SDS in PBS, diluted to a final detergent concentration of 0.4% SDS, 0.6% NP-40 in PBS. Streptavidin enrichment and trypsin digestion was conducted on the eluate for label free quantification as described above.

Neuronal Culture

Mouse breeding is essentially as described previously (Ref Samowitz et al. 2025). Forebrains containing cortex and hippocampal formation were dissected from postnatal day 0 (PND 0) mouse pups to isolate primary cortical neurons in dissection media (culture grade $H_2O$ (Fisher Scientific: SH3052902), 10% 10x HBSS without $Ca^{2+}$ and $Mg^{2+}$ (Invitrogen: 14185052), 2% HEPES (Invitrogen: 15630080), 1% pyruvate (Invitrogen: 11360070), 1% glucose solution (Thermo: A2494001), and 0.02% Gentamicin (Invitrogen: 15710064). The cortices were placed in a digestion solution containing dissection media and 20 active units/mL of papain (Worthington: LS003124) for 30 min at 37 °C. Tissues were washed and triturated in plating medium consisting of Neurobasal (Invitrogen: 21103049) containing 5% FBS, heat inactivated (Invitrogen: 10082139), 2% Glutamax-I, (Invitrogen: 35050061), and 0.02% Gentamicin (Invitrogen: 15710064). Cells were then centrifuged for 5 min at 800xg and resuspended in plating medium at 300 µL per brain. Cell suspension was then diluted into feeding medium consisting of Neurobasal-A (Invitrogen: 10888022), 2% Glutamax-I, and 0.02% Gentamicin, 2% B-27 supplement (Invitrogen: 17504044),

Neuronal RNA Seq

Neurons were isolated from *Syngap1* conditional rescue (*Syngap1*$^{+/ls}$) mice and plated using a BioTek EL406 (Agilent Technologies) into 384-well plates pre-coated with poly-D-lysine (PDL) (Aurora ABE2-01200B-PDL) at 15,000 cells in 80 µL/ well and placed in 37 °C incubator (*42*). A solution of 1% agarose was placed in the evaporation border wells prior to plating to minimize edge effects. On DIV 3, 10 µM 5-fluoro-2′-deoxyuridine (+FUDR) was added cultures to suppress the proliferation of glia. For 2-week treatments of SR-1815, cultures were treated with vehicle (DMSO) or SR-1815 at 1.5 µM on DIV 0, 3, 7, and 10 by replacing 50% of the conditioned media with fresh feeding media (+FUDR). For 4 h treatments, 50% of media was replaced with fresh feeding media on DIV 3, 7, and 10. On DIV 14 cultures were supplied vehicle or SR-1815 at 4.4 µM for 4 h through 50% media exchange. On DIV 14 media was removed and 20 µL of DNA/RNA Shield (Zymo: R1100-50) was added to each well. Sample was collected and RNA was extracted using Zymo Quick-RNA MicroPrep kit (Zymo: R1050) following manufacturer protocol.

<u>Neuronal Proteomics:</u>

Neurons were isolated from *Syngap1*$^{+/ls}$ mice and cell suspension was plated into PDL coated 24-well plates at 250,000 cells / well. Neurons were treated with vehicle or SR-1815 at 1.5 µM on DIV 0, 3, 7, and 10 by 50% media exchange (+FUDR at DIV 3-14). On DIV 14, wells were washed 2X with PBS and lysed in a buffer consisting of 2% SDS, 50 mM HEPES pH 8.5, protease inhibitor (Roche:05892791001), and phosphatase inhibitor (Roche: 04906845001). Lysates were generated and processed as previously described in "label free nuclear proteomics".

<u>Neuronal Western Blot</u>

Neurons were prepared and treated as indicated in neuronal proteomics. On DIV 14, plates were washed with PBS twice and proteins were extracted by sonication in a buffer consisting of 2% SDS, 50 mM Sodium Borate, 1X Halt Protease and Phosphatase inhibitors. Sample protein concentrations were measured using Pierce BCA Protein Assay Kit (Thermo: 23225) and adjusted to normalize protein content. 10 µg of protein per sample was loaded and separated by SDS-PAGE on 10% Criterion TGX Stain-Free gels (BioRad: 5678035), transferred to low fluorescence PVDF membranes (45 µm pore size) (Amersham: GE10600004) with the Trans-Blot Turbo System (BioRad). Membranes were imaged for total protein using BioRad ChemiDoc imaging system, blocked with 1% BSA-TBS-T for 1 h, and then probed with primary antibodies at 4 °C overnight. Membranes were washed 3X with TBS-T, incubated with secondary antibodies, washed, and imaged for chemiluminescence.

<u>Neuronal Morphology and Spine Quantification</u>

Neurons were isolated from *Syngap1* constitutive (*Syngap1*$^{+/-}$) mice and plated into 384-well plates at 15,000 neurons per well containing vehicle or SR-1815 at 1.5 µM (*43*). On DIV 3 neurons were transduced with AAV9 vectors AAV pCAG-FLEX-EGFP-WPRE (Addgene: 51502-AAV9) (MOI=100,000 vp/cell) and pENN.AAV.hSyn.Cre.WPRE.hGH (AAV9) (Addgene: 105553-AAV9) (MOI=10 vp/cell) by 50% media exchange with vehicle or SR-1815 (+FUDR). Neurons were given 50% media exchanged containing vehicle or SR-1815 on DIV 7 and 10 (+FUDR at DIV 7-14). Imaging was performed on DIV 14 using an IN Cell 6000 (20x magnification). Tracing data were obtained by imaging the entire well and stitched together using ImageJ. Dendritic morphology was traced using Fiji-ImageJ plugin Simple Neurite Tracer (SNT) (PMID:21727141). A sample of 32 neurons per group were randomly selected and traced blinded. Data represent the average branch length in microns for all dendrites of the traced neuron. For spine density analysis,

2-3 dendrites were selected per traced neuron and spines were counted using SNT and normalized to dendritic length. Spine intensity was normalized to dendritic intensity.

Immunocytochemistry

Neurons were isolated from *Syngap1* constitutive (Syngap1$^{+/-}$) mice and plated onto PDL coated glass coverslips (Neuvitro: GG-18-15H) into 12-well plates at 500,000 neurons per well containing vehicle or SR-1815 at 1.5 µM, 3 coverslips per condition (*43*). A 50% media exchange occurred on DIV 3, 7, and 10 containing vehicle or SR-1815 (+FUDR at DIV 3-14). At DIV 14 coverslips were transferred to a new plate containing a fixation solution of 4% PFA, 4% Sucrose, 80, mM PIPES, 2 mM $MgCl_2$, and 5 mM EGTA pH 6.8 for 10 min. Coverslips were washed 3X with PBS and incubated in blocking solution of 2% BSA and 0.1% Triton-X in PBS for 1 h. Coverslips were incubated blocking solution containing MAP2 primary antibody (1:1000) (SySy: 188004) overnight at 4 °C and washed 3X with PBS. Goat anti-Guinea pig Alexa Fluor 647 Secondary antibody (10 µg/mL) (Invtirogen: A-21450) was added to blocking buffer and incubated for 1 h at room temperature. Coverslips were washed 3X with PBS and mounted on to slides with VECTASHIELD HardSet Antifade Mounting Medium with DAPI (VectorLabs: H-1500) for 24 h. Images were taken using an IN Cell 6000 from 36 randomly selected fields (60X magnification) per coverslip. Mean pixel intensity was calculated for each image using Fiji-Image J.

Syngap1 Assay

Neurons were isolated from *Rosa26*$^{+/fLuc}$; *Syngap1*$^{hb/fl}$ mice (Ref: Samowitz et al. 2025) were transduced with 30,000 viral particles/cell of pENN.AAV.hSyn.Cre.WPRE.hGH (AAV9) (Addgene: 105553-AAV9) to induce haploinsufficiency and plated into 384-well plates at 10,000 cells / well and administered compounds with a 25 nL pin tool (V&P Scientific) with vehicle, SR-1815-Dz, selected inhibitors, and SR-1815 (+FUDR at DIV 0). A 50% media exchange was performed on DIV 3, 7, and 10 (+FUDR at DIV 3-14) and subsequently pinned with vehicle or compounds in an 8pt-dose response. Vehicle or TRKB-FC Chimera was dissolved into media and administered using a 50% media exchange using Certus Flex dispenser (Trajan Scientific and Medical). At DIV 14 a Dual-Luciferase Reporter assay was performed for SynGAP protein expression analysis as previously described (Samowitz et al. 2025). Data shown for each inhibitor shown was selected showing the highest % normalized SynGAP expression before the onset of compound toxicity (Normalized fLuc and nBiT signals >1). % Normalized SynGAP expression was calculated for each sample well as follows: (2^($log_2$(Lum nBiT Sample)-mean($log_2$(Lum nBiT Vehicle Ctrl))-($log_2$(Lum fLuc Sample)-mean($log_2$(Lum fLuc Vehicle Ctrl)))) -1)*100.

KINOME*scan*™

We contracted Eurofins DiscoverX (Fremont, CA) to perform the scanMAX assay within the larger KINOME*scan*™ screening platform. KINOME*scan*™ assays do not require ATP and thereby report true thermodynamic interaction affinities, as opposed to IC50 values, which can depend on the ATP concentration. The largest commercial kinase panel available, scanMAX contains a set of 468 kinases covering AGC, CAMK, CMGC, CK1, STE, TK, TKL, lipid and atypical kinase families, plus important mutant forms. scanMAX is an ideal choice for all stages of drug discovery and development, from lead discovery & hit identification to lead optimization, preclinical and compound safety. Kinome-wide annotation of compound selectivity enables informed decisions about therapeutic opportunities and potential off-target liabilities which could otherwise be missed in smaller kinase panels. This panel includes: 453 human kinases, 3 pathogen kinases, 56 disease relevant mutants, 132 tyrosine kinase assays, and 20 lipid kinases.

Kinase assays

For most assays, kinase-tagged T7 phage strains were grown in parallel in 24-well blocks in an E. coli host derived from the BL21 strain. E. coli were grown to log-phase and infected with T7 phage from a frozen stock (multiplicity of infection = 0.4) and incubated with shaking at 32 °C until lysis (90-150 min). The lysates were centrifuged (6,000 x g) and filtered (0.2 µm) to remove cell debris. The remaining kinases were produced in HEK-293 cells and subsequently tagged with DNA for qPCR detection. Streptavidin-coated magnetic beads were treated with biotinylated small molecule ligands for 30 min at room temperature to generate affinity resins for kinase assays. The liganded beads were blocked with excess biotin and washed with blocking buffer (SeaBlock (Pierce), 1 % BSA, 0.05 % Tween 20, 1 mM DTT) to remove unbound ligand and to reduce non-specific phage binding. Binding reactions were assembled by combining kinases, liganded affinity beads, and test compounds in 1x binding buffer (20 % SeaBlock, 0.17x PBS, 0.05 % Tween 20, 6 mM DTT). Test compounds were prepared as 100x stocks in 100% DMSO and directly diluted into the assay. All reactions were performed in polypropylene 384-well plates in a final volume of 0.02 mL. The assay plates were incubated at room temperature with shaking for 1 h and the affinity beads were washed with wash buffer (1x PBS, 0.05 % Tween 20). The beads were then re-suspended in elution buffer (1x PBS, 0.05 % Tween 20, 0.5 µM non-biotinylated affinity ligand) and incubated at room temperature with shaking for 30 min. The kinase concentration in the eluates was measured by qPCR.

Calculation of activity (%Ctrl)

SR1815 was screened at 1mM, and results for primary screen binding interactions are reported as '% Ctrl', where lower numbers indicate stronger hits in the matrix **(Table S22)**.

%Ctrl Calculation

$$\left(\frac{\text{test compound signal - positive control signal}}{\text{negative control signal - positive control signal}}\right) \times 100$$

test compound = SR1815
negative control = DMSO (100%Ctrl)
positive control = control compound (0%Ctrl)

Selectivity Score (S-scores)

Selectivity Score or S-score is a quantitative measure of compound selectivity. It is calculated by dividing the number of kinases that compounds bind to by the total number of distinct kinases tested, excluding mutant variants.

S = Number of hits / Number of assays

This value can be calculated using %Ctrl as a potency threshold (below) and provides a quantitative method of describing compound selectivity to facilitate comparison of different compounds.

S(35) = (number of non-mutant kinases with %Ctrl <35)/(number of non-mutant kinases tested)

S(10) = (number of non-mutant kinases with %Ctrl <10)/(number of non-mutant kinases tested)

S(1) = (number of non-mutant kinases with %Ctrl <1)/(number of non-mutant kinases tested)

Kinase Dendrograms

Images were generated using TREEspot™ Software Tool and are reprinted with permission from KINOMEscan®, a division of DiscoveRx Corporation, © DISCOVERX CORPORATION 2010. Dendrograms for Sunitinib, Imatinib, and Erlotinib were provide as reference data by DiscoveRx Corporation and can be found at https://www.eurofinsdiscovery.com/StudyManagement/Treespot.

Experiments in Cancer Lines

MDA-MB-468 (ATCC HTB-132), A549 (ATCC CRM-CCL-185), MCF7 (ATCC HTB-22), and MIA PaCa-2 (ATCC CRL-1420) cell lines were cultured according to the protocols provided by ATCC at 37 ºC and 5% $CO_2$ in a Forma™ Steri-Cycle™ $CO_2$ Incubator (cat. 370, Thermo Fisher Scientific). Reaching about ~70% confluency, cells grown in 75 $cm^2$ flasks (cat. 430641U, Corning, Corning, NY) were washed twice with 10 mL of DPBS (cat. 14190250, Life Technologies, Carlsbad, CA) and dissociated by adding 2 mL Trypsin-EDTA (0.25%) solution (cat. 25200056, Thermo Fisher Scientific) followed by incubation at 37 ºC for 10 min. Further cell dissociation was facilitated by gently pipetting the suspension up and down several times until no cell aggregates were observed by visual inspection under a stereomicroscope. Following the addition of fresh culture medium (8 mL), the cell density was determined by counting the cells in a hemocytometer. Cells were diluted and plated onto flat bottom, 96-well cell culture plates (cat. BD353377, Thermo Fisher Scientific) by transferring 100 µL cell suspension to each well using a multichannel pipette at a final surface densities of 2000, 1000, 700, and 1000 cells/well, for MDA-MB-468, A549, MCF7, and MIA PaCa-2 cells, respectively.

After 24 h of incubation at 37°C and 5% $CO_2$, cells were treated with SR-1815, or sunitinib. The compounds were first dissolved in dimethyl sulfoxide (DMSO, cat. D2650, Sigma-Aldrich) at a concentration of 4 mM. Second, twelve-step serial 1:2 dilutions of compound solutions were prepared in DMSO using solvent-resistant polypropylene microplates (cat. 3357, Corning). Third, compound plates were prepared by adding 0.7 µL of DMSO-based compound solutions to each well of a 96 well plate containing 139.3 µL of culture media using a multichannel pipette (resulting in a 200-fold dilution). Wells in the first and last rows were used exclusively for negative and positive controls, respectively. In these rows, 0.7 µL of DMSO was combined with 139.3 µL of culture media. Solutions were mixed by shaking the compound plate for 1 min at room temperature at 800 rpm using a microplate shaker (cat. 12620-926, VWR). Finally, 100 µL of diluted compound solutions were transferred from the compound plate to the assay plate (containing cell cultures in 100 µL of culture media) using a multichannel pipette (resulting in an extra 2-fold dilution). NOTE: The DMSO concentration in the assay plate was 0.25%. The final assay concentration of compounds was between 10 µM and 5 nM. All measurements were carried out in triplicate. Cells treated with pure DMSO were used as the negative control, while wells with culture medium only (no cell suspension) were used as the positive control. A plate layout map is shown in Figure 1.

After 3 or 5 days of incubation at 37 °C and 5% $CO_2$, assay plates were equilibrated to room temperature. The cell culture medium was removed from the plates, and 100 μL of a room-temperature mixture of 1:1 DPBS and CellTiter-Glo® 2.0 Reagent (cat. G9242, Promega) was dispensed into each well using a multichannel pipette. Plates were shaken for 2 min at 800 rpm and incubated at room temperature for an additional 10 min. Luminescence was recorded using a CLARIOstar Plus microplate reader (BMG LABTECH) with an integration time of 0.8 sec per well. The luminescence values against the compound concentration were plotted using OriginPro 2017 software (OriginLab), and the half maximal effective concentration ($EC_{50}$) was determined by fitting the 12-point dose-response data to the Hill equation:

Eq. (1) $Y = Y_{min} + (Y_{max} - Y_{min})\left(1 - \frac{1}{1+\left(\frac{c_{compound}}{EC_{50}}\right)^{Hill}}\right)$, where $Y$ is the luminescence signal, $c_{compound}$ is the compound concentration, $Y_{min}$ and $Y_{max}$ are the two asymptotes, and *Hill* is the Hill-constant.

### Supplementary Figures

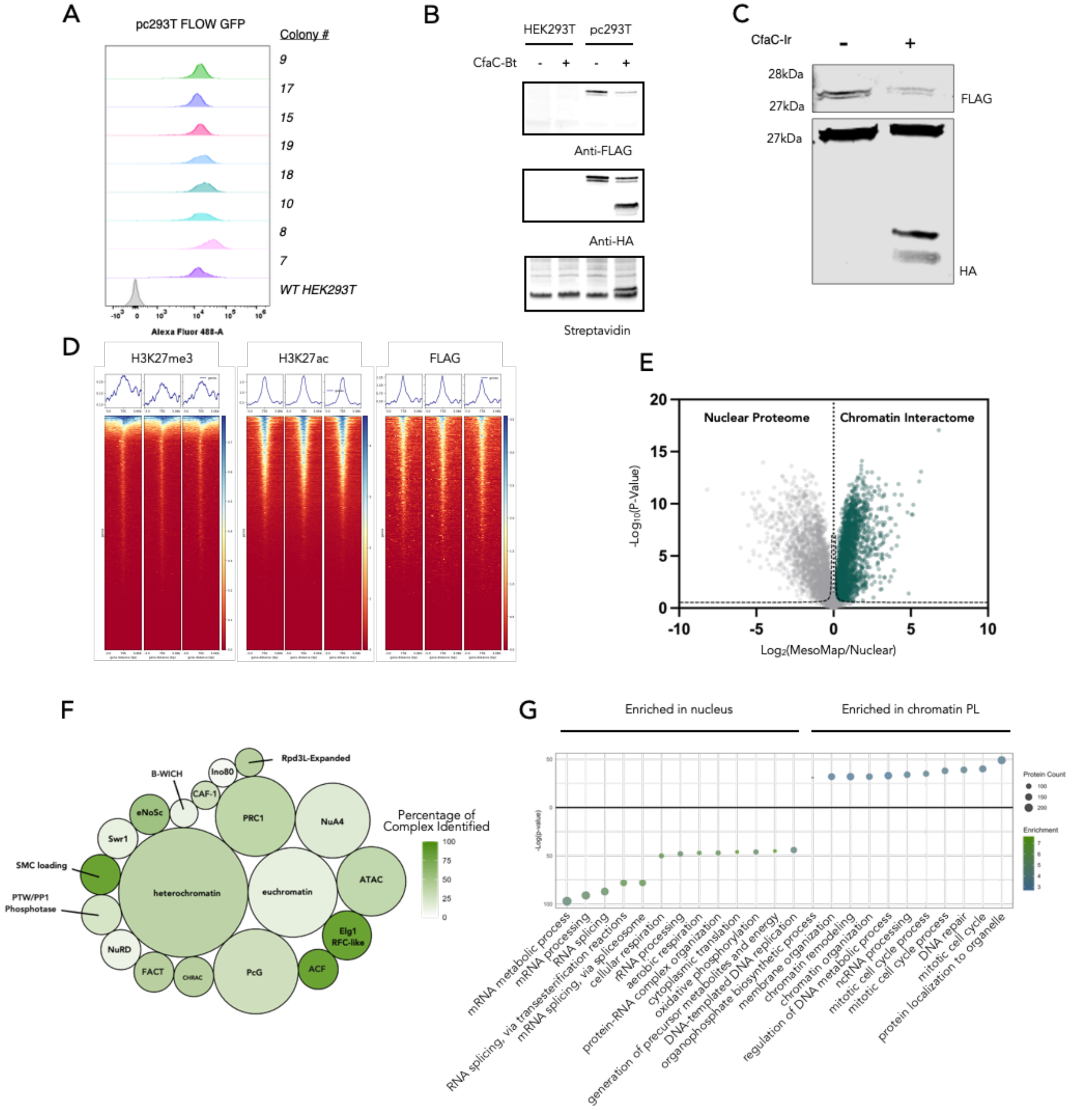


**Fig. S1. MesoMap Development.** (**A**) Flow cytometry of GFP expression in pc293T single cell colonies. (**B**) Western blotting of CfaC-Biotin splicing onto histone constructs in pc293T cells. (**C**) Western blotting of CfaC-Ir splicing onto histone constructs in pc293T cells. (**D**) Heatmaps of CUT&Tag of pc293T cells showing H3K27me3, H3K27ac, and FLAG peaks centered on transcription start site +/- 3kb. Reads normalize to counts per million. Generated using deepTools. (**E**) Volcano plot of nuclear proteome vs. MesoMap proteome. Green points are all proteins that were significantly enriched at chromatin. (**F**) Bubble plot of chromatin remodeling complexes.

Data included is the significant hits enriched by MesoMap in E. Size of bubble correlates to number of members in complex and color represents percentage of members identified in chromatin proximity labeling. (**G**) Gene ontology of the hits significantly enriched in nucleus and at chromatin.

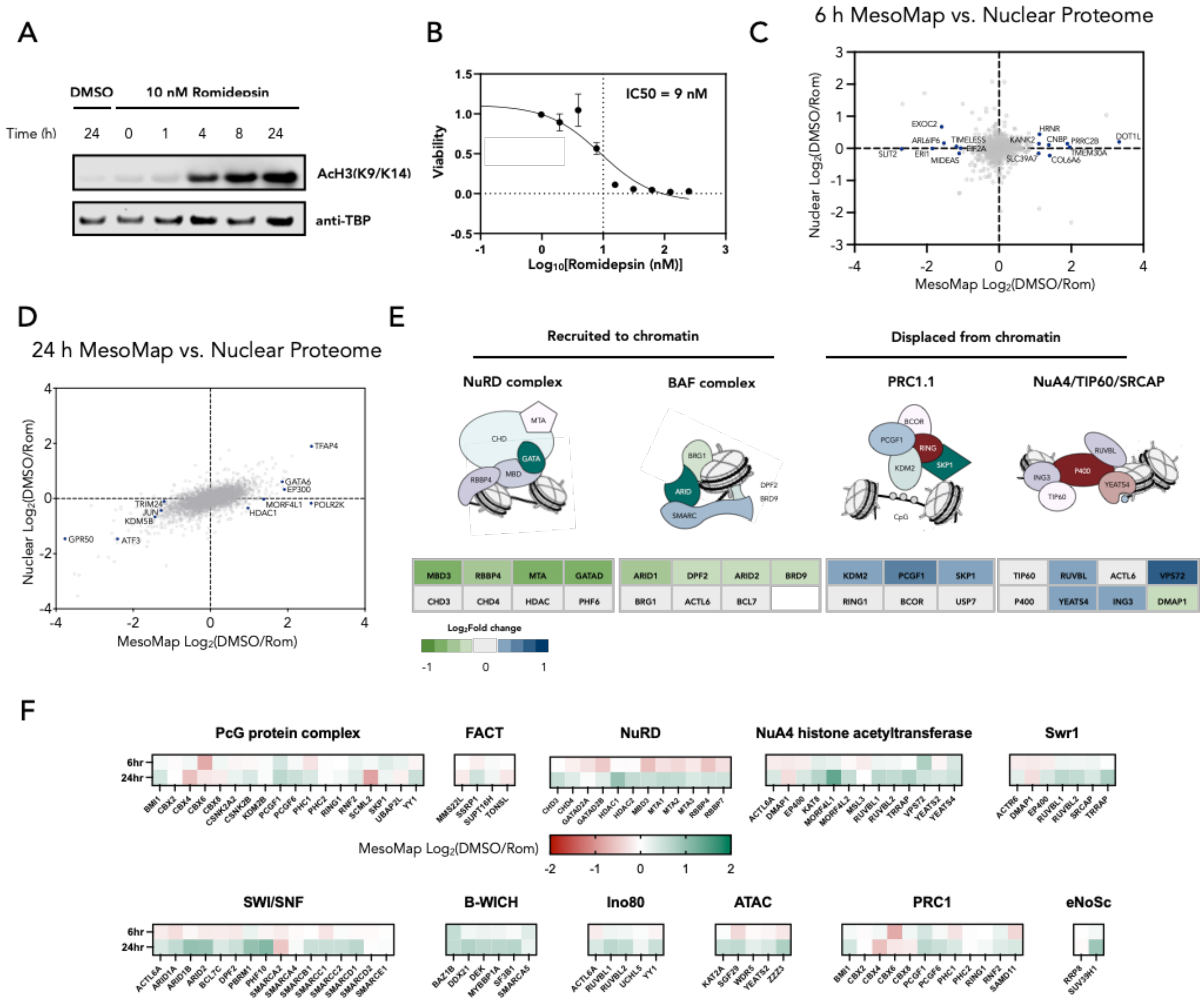


**Fig. S2. MesoMap exploration of romidepsin impact on chromatin remodelers.** (**A**) Western blotting for H3K9/K14 acetylation in pc293T cells treated with 10 nM romidepsin. (**B**) Cell viability assay of pc293T cells treated for 72 h with indicated concentration of romidepsin. Values normalized to DMSO. (**C**-**D**) Scatter plots of MesoMap (x-axis) and nuclear proteome (y-axis) changes in response to romidepsin treatment. Genes of interest are labeled and highlighted in blue. (**E**) MesoMap data for select chromatin remodeling complexes at 24 h treatment with 10 nM romidepsin. Illustrations of complex (above) and heatmap of member proteins $Log_2$(DMSO/romidepsin) MesoMap data. (**F**) MesoMap data of select chromatin remodeling complexes at 6 h and 24 h treatment points with 10 nM romidepsin. Values in the heatmap are $Log_2$(DMSO/romidepsin) of MesoMap data.

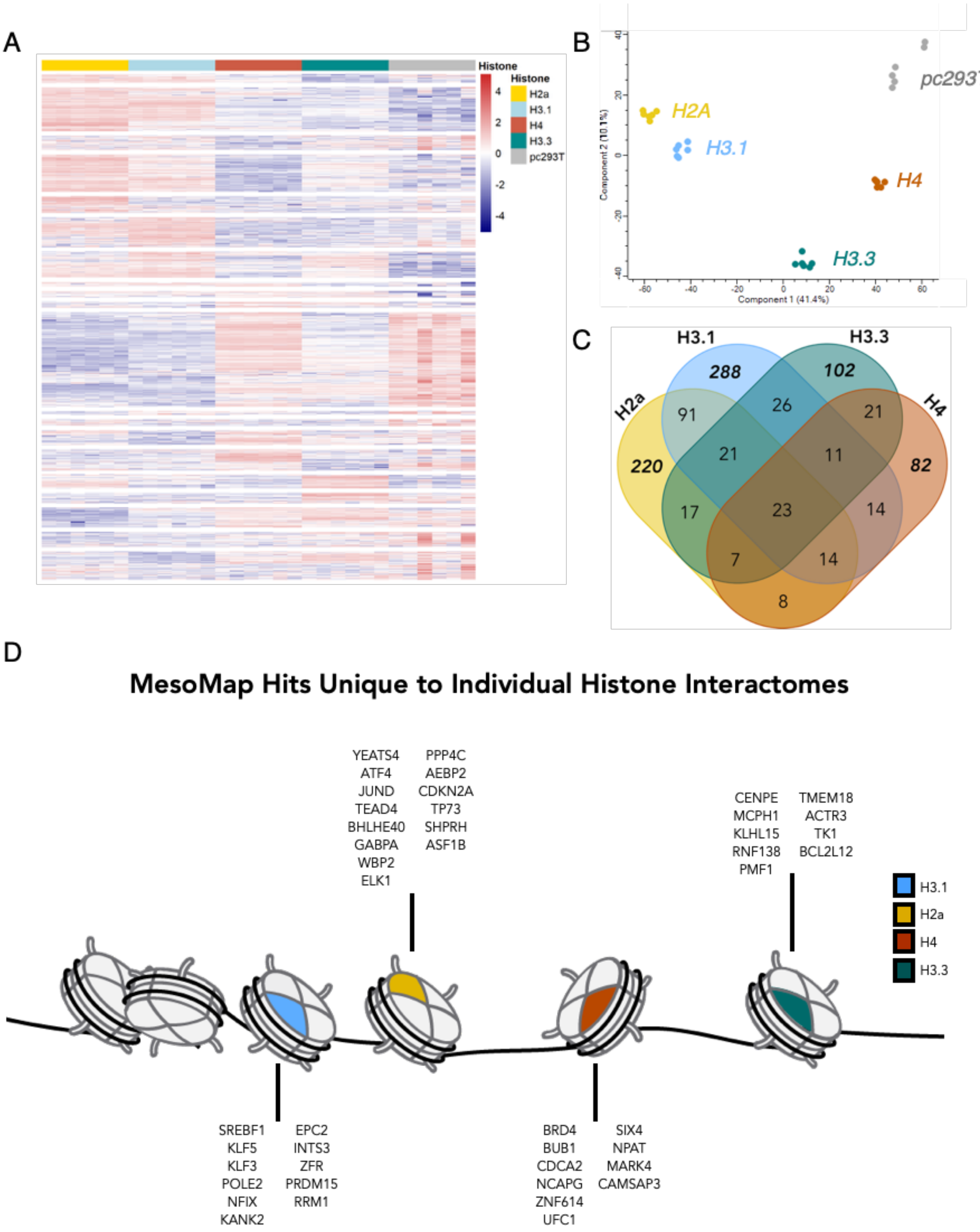


**Fig. S3. MesoMap captures interactions across all of chromatin, which cannot be reproduced with single histone proximity labeling.** (**A**) Heatmap of $Log_2$ normalized abundance of proteins enriched by µMap for each histone construct as well as pc293T cells. (**B**) Principal component analysis of enriched data of individual histone constructs as well as pc293T cells. (**C**) Venn diagram of significant differentially chromatin associated genes in response to 24 h treatment of

10 nM romidepsin in μMap experiments of individual histones. (**D**) Mesomap captures aggregate chromatin interactions that cannot be replicated with single histone construct. Highlighted proteins are significant differentially enriched proteins from MesoMap that only met significance in one histone μMap experiment. MesoMap and μMap differentially enriched genes are from 24 h 10 nM romidepsin treatment experiments.

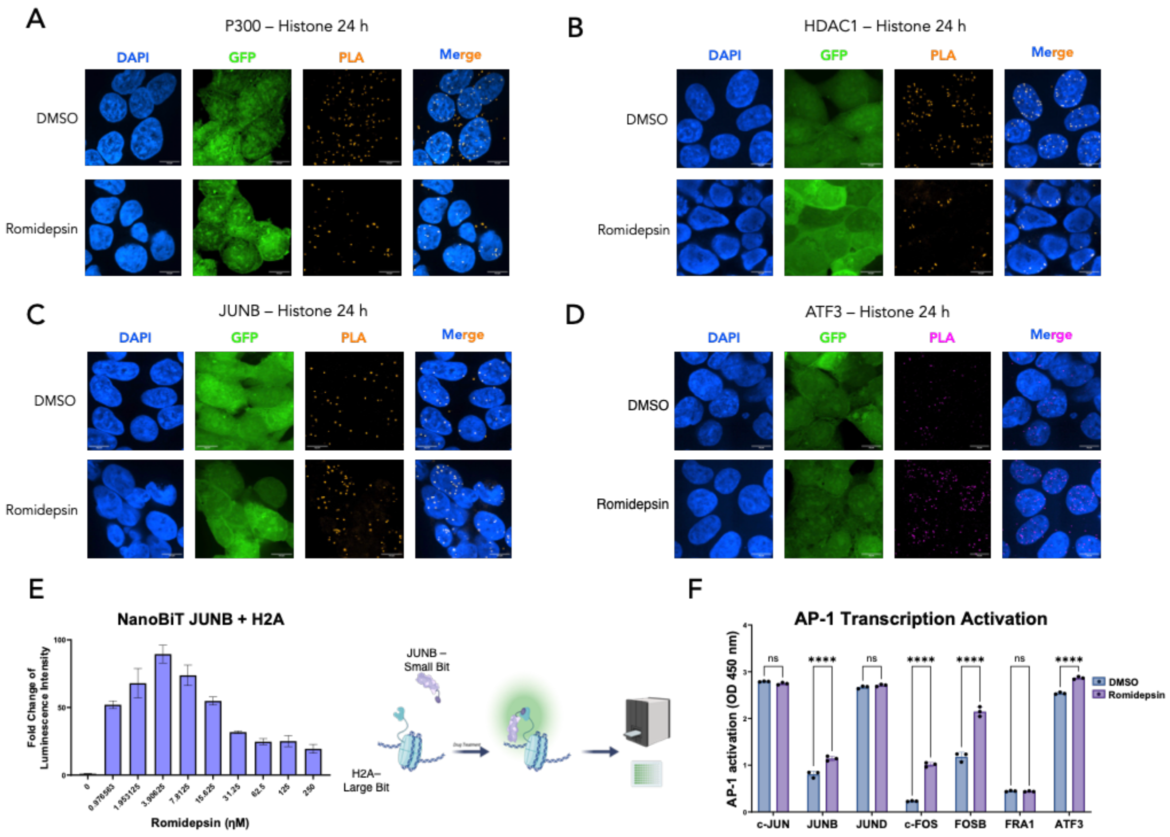


**Fig. S4. Concurrent MesoMap and phosphoproteomic analysis of transcriptional regulators at chromatin.** (**A-D**) Proximity ligation assay representative images for quantification shown in figure 2K. (**E**) Nanobit results detecting JUNB and Histone H2A protein interactions at 24 h with treatment of indicated concentration of romidepsin. (**F**) ELISA based DNA binding assay of AP-1 family members to the CRE motif. Cells were treated with 10 nM romidespin or DMSO for 24 h, lysed, and lysate used for ELISA.

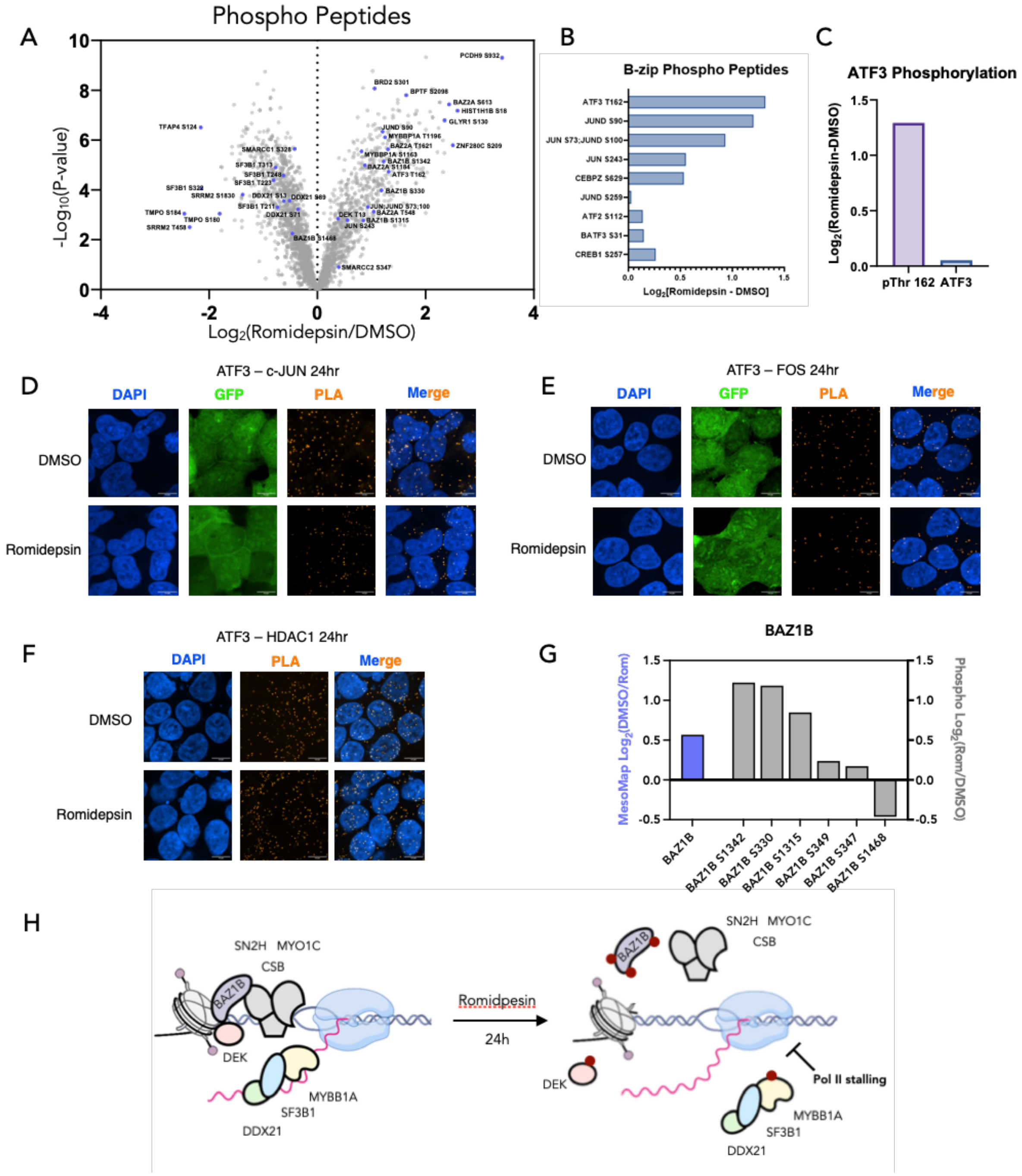


**Fig. S5. Romidepsin impact on phosphorylation state, bZip cistrome, and B-WICH chromatin interactions.** (**A**) Volcano plot of phospho peptides identified in phosphoproteomic analysis of cells treated with DMSO or romidepsin for 24h. (**B**) Bar plot of select B-zip phosphopeptides differentially phosphorylated in pc293T cells treated with romidepsin. (**C**) Bar graph of ATF3 pThr 162 and global ATF3 expression changes in response to romidepsin treatment. (**D-F**) Proximity ligation assay representative images for quantification shown in figure 2M. (**G**) Bar graph of global BAZ1B expression change (Blue) and specific phosphosite changes (grey) in response to romidepsin treatment. (**H**) Illustration of phosphorylation events of B-WICH complex members, leading to dissociation from chromatin and Pol II stalling.

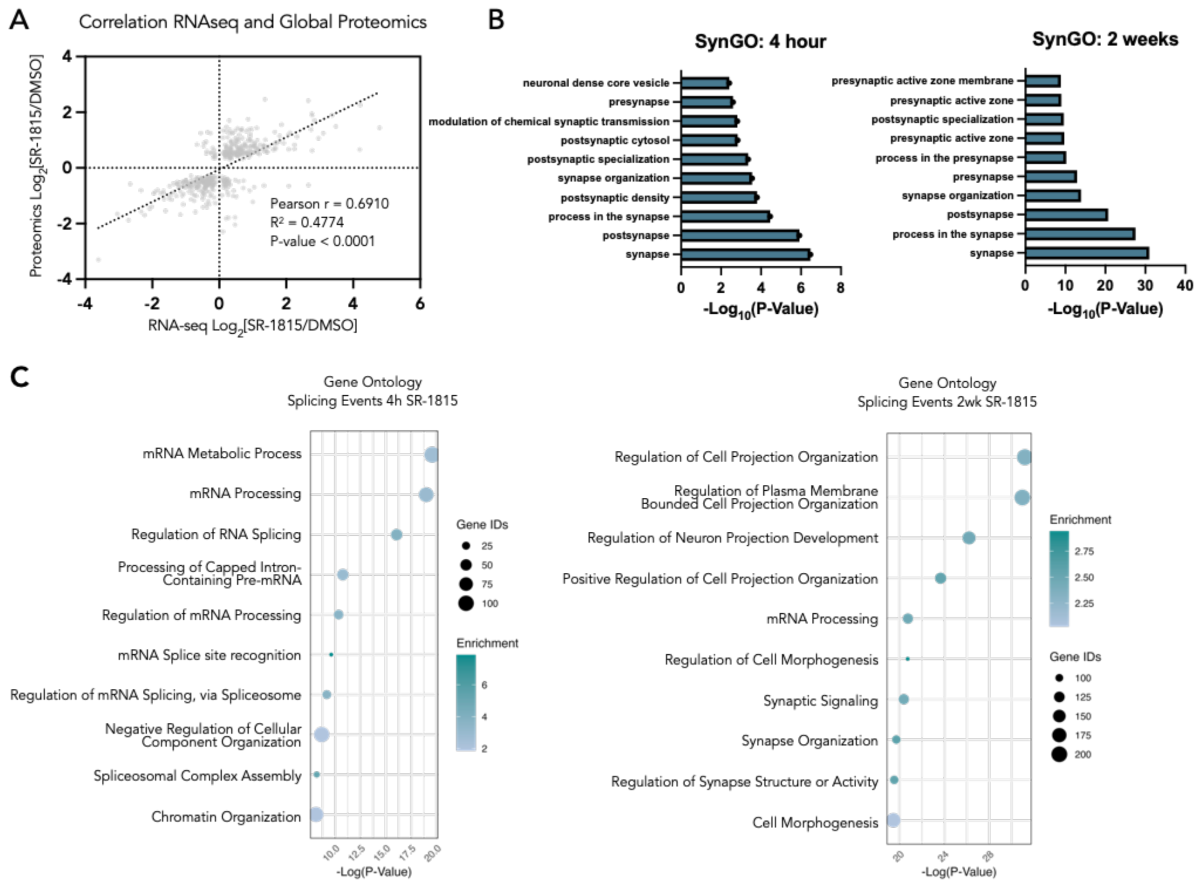


**Fig. S6. RNAseq in SR-1815 treated neurons shows differential splicing activity.** (**A**) Scatterplot showing Pearson correlation of hits that met significance in global proteomics and RNA-seq datasets in neurons treated with SR-1815 for 2 wk. (**B**) SynGO of RNA-seq data from neurons treated with SR-1815 for indicated time point. (**C**) Gene ontology of significant splicing events in neurons treated with SR-1815 at indicated duration.

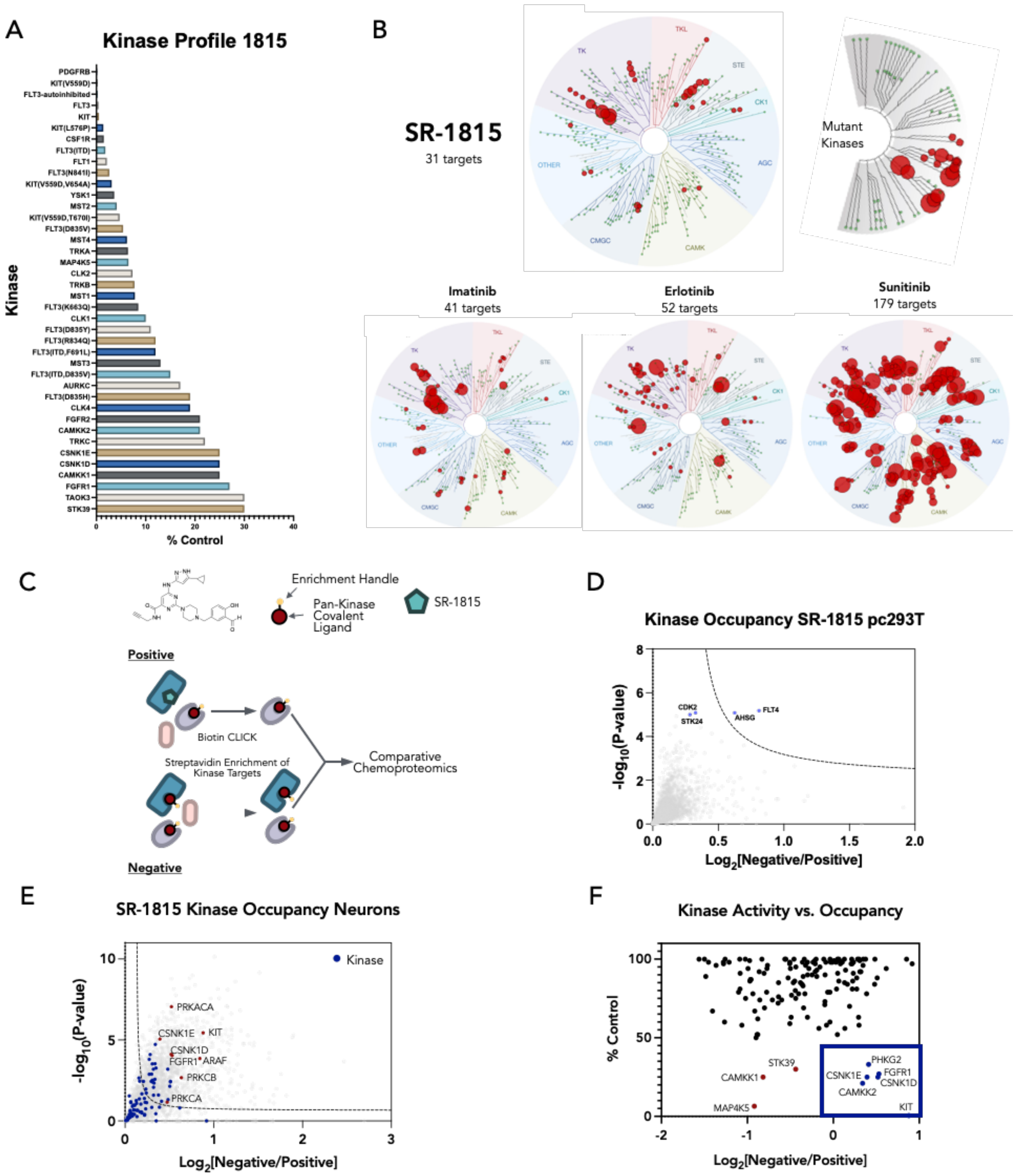


**Fig. S7. SR-1815 kinomeSCAN results and evidence that it engages kinases in neurons. (A)** Bar graph of hits from kinome screen of SR-1815. (**B**) Kinase profiles of SR-1815 and FDA approved kinase inhibitors. (**C**) Diagram of in cell kinase occupancy experiment. (**D**) Volcano plot of kinase occupancy of SR-1815 in pc293T cells. (**E**) Kinase occupancy data of SR-1815 in neurons. (**F**) Scatterplot of kinome screen data of inhibition reported as % Control activity (y-axis) versus the kinase occupancy $log_2FC$ in neurons (x-axis). Proteins highlighted in lower right are kinases that demonstrate SR-1815 binding as well as inhibition.

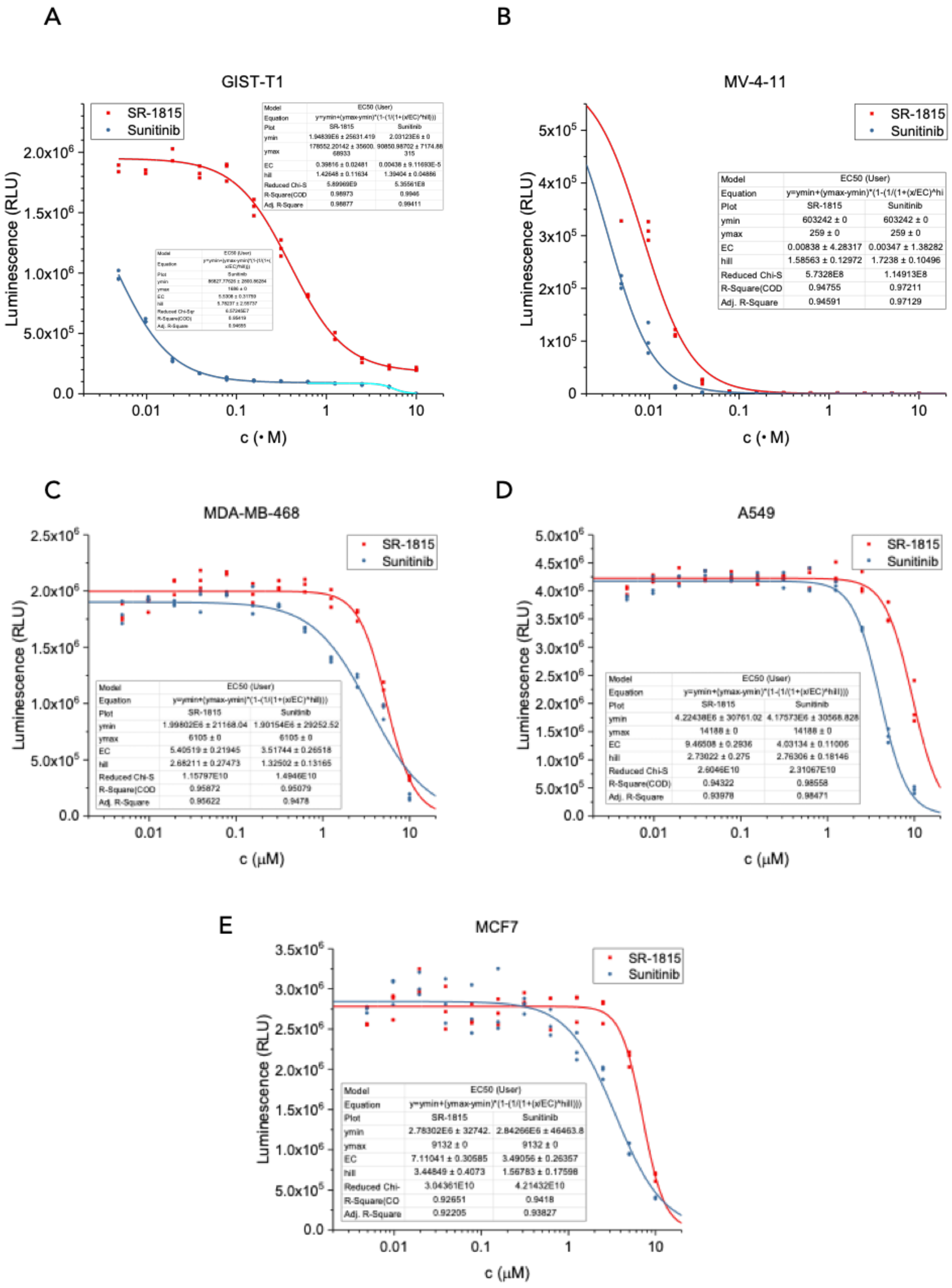


**Fig. S8**. **SR-1815 is a highly potent antiproliferative agent in cancer models.** (**A-E**) Cell viability assay of SR-1815 and Sunitinib in cancer cell lines: MV-4-11 (acute myeloid leukemia) and GIST-T1 (gastrointestinal stromal tumor), MDA-MB-468 (metastatic adenocarcinoma of the breast), A549 (lung carcinoma), and MCF7 (adenocarcinoma of the breast).
